# Collagen IV of basement membrane: V. Bromide-mediated sulfilimine bonds interlock the quaternary structure of NC1-hexamer of scaffolds enabling metazoan evolution

**DOI:** 10.64898/2026.02.20.707065

**Authors:** Bradley P. Clarke, Vadim Pedchenko, Tetyana Pedchenko, Monica Moran, Jacob Edwards, Kyle Vallone, Carl Darris, Gautam Bhave, Patrick Page-McCaw, Julie K. Hudson, Sergei P. Boudko, Billy G. Hudson

**Author notes:** co-first authors (Bradley P. Clarke, Vadim Pedchenko, Tetyana Pedchenko). Co-senior Author (Bradley P. Clarke, Billy G. Hudson). Aspirnauts Aspirnaut is a K-20 Science, Technology, Engineering, and Math (STEM) pipeline for increasing the diversity and wellness of the STEM workforce. The holistic training approach features guided discovery science that is augmented with guided professional skills development, guided self-discovery, and wellness training. High school students from rural America and diverse backgrounds engage in hands-on discovery science for 6 weeks while in residence at Vanderbilt University Medical Center, and diverse undergraduate students engage for 10 weeks. The experience provides students with the tools and empowerment to effect positive change in themselves, their families, and communities for generations to come.

## Abstract

Collagen-IV (Col-IV) scaffolds, a primordial basement membrane component, enabled animal multicellularity, evolution and adaptation. These scaffolds provide tensile strength and tether macromolecules, forming supramolecular complexes that interact with cell-surface receptors and influence cell-behavior. Triple-helical Col-IV protomers, composed of three α-chains, with a trimeric globular NC1-domain at the C-terminus, oligomerize forming a NC1-hexamer structure that connects adjoining protomers of Col-IV^α121^, Col-IV^α556—α121^, and Col-IV**^α345^** scaffolds. Hexamer formation and stability are driven by the extracellular chloride concentration-“chloride pressure”. Hexamer structure is reinforced by six sulfilimine bonds forming covalent crosslinks that weld together trimeric NC1-domains of adjoining protomers. We recently found evidence that sulfilimine bonds, independent of chloride, stabilize the quaternary structure of the Col-IV**^α345^** hexamer of the Col-IV**^α345^** scaffold. Here, we sought to determine whether this function also pertains to the Col-IV**^α121^** scaffold that occurs ubiquitously across the animal kingdom, and whether bromine, a cofactor of peroxidasin in bond formation, are evolutionary conserved. We found that sulfilimine bonds stabilized the quaternary structure of the Col-IV**^α121^** hexamer of bovine, mouse and a basal cnidarian, *Nematostella vectensis*, and that the mechanism of bond formation mediated by peroxidasin and bromide is evolutionary conserved. Analyses of the crystal structure of the NC1-hexamer revealed that sulfilimine bonds covalently fasten a clasp-motif across the trimer-trimer interface, interlocking the domain-swapping region of neighboring subunits, which reinforces the hexamer quaternary structure imposed by chloride conformational constraints. Collectively, our findings reveal that the sulfilimine-bond reinforcement is a critical event in Col-IV scaffold assembly enabling multicellularity, evolution and adaptation of metazoans, beginning with ancient cnidarians.

## Introduction

The collagen IV (Col-IV) scaffold, a primordial innovation, enabled the assembly of a fundamental architectural unit of epithelial tissues - a basement membrane (BM) attached to polarized cells (Fidler et al., 2017, 2018; P. S. Page-McCaw et al., 2025). The BM is a specialized form of extracellular matrix with numerous functions ranging from tissue architecture to orchestrating cell behavior and ultrafiltration (Brown et al., 2017; Clay & Sherwood, 2015; McCall et al., 2014; A. Page-McCaw & Ferrell, 2025; Pozzi et al., 2017). Col-IV is a family of six collagen IV α-chains (α1 to α6) that self-assemble inside of the cell into three triple-helical protomers of distinct α-chain compositions (α121, α345 and α565) (Hudson et al., 2003; Khoshnoodi et al., 2008; Yurchenco & Ruben, 1987). On the outside of cells, protomers self-assemble into three supramolecular scaffolds, noted as Col-IV^α121^, Col-IV^α345^, and Col-IV^α556—α121^. Scaffolds confer tensile strength to tissues and tether diverse macromolecules, including laminin, nidogen, proteoglycan, and growth factors, forming supramolecular complexes that interact with cell surface receptors (Brown et al., 2017; Fidler et al., 2018). The Col4^α121^ scaffold is a ubiquitous component of metazoan BMs, whereas Col-IV^α345^ and Col-IV^α556—α121^ are specialized components of various vertebrate tissues (P. S. Page-McCaw et al., 2025).

During scaffold assembly, Col-IV protomers oligomerize head-to-head via trimeric globular NC1-domains at the C-terminus, forming a NC1-hexamer structure at the trimer-trimer interface (Boudko et al., 2021; Casino et al., 2018; Sundaramoorthy et al., 2002; Than et al., 2002.). Hexamer formation and stability of the quaternary structure is driven by the extracellular chloride concentration - “chloride pressure” (Fig. 1 first arrow) (Boudko et al., 2023; Cummings et al., 2016; Pedchenko et al., 2019). Hexamer structure is further stabilized by six sulfilimine bonds forming covalent crosslinks that weld together trimeric NC1-domains of adjoining protomers (Fig. 1 second arrow) (Boudko et al., 2021; McCall et al., 2014; R. Vanacore et al., 2009). Sulfilimine bond formation uniquely involves bromide as cofactor for peroxidasin (Bhave et al., 2012; McCall et al., 2014). Prior to this finding, Br-was considered a trace ion without a biological function. Numerous in vitro studies have shown that peroxdasin catalyzes sulfilimine bond formation and utilizes HOBr as the oxidant (Bhave et al., 2012; McCall et al., 2014; Ronsein et al., 2014; Roy & Gauld, 2023).

**Figure 1.**
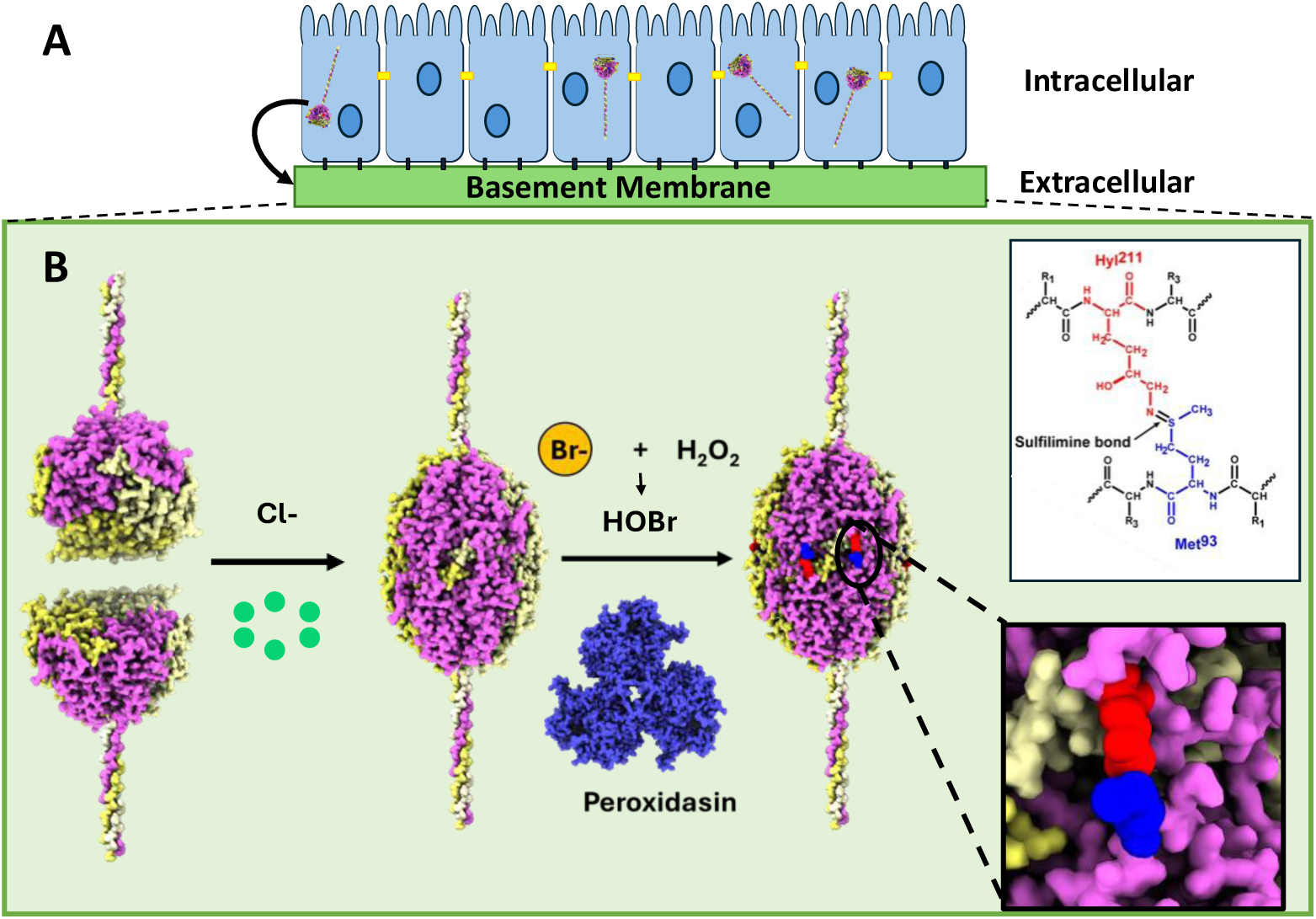
The cooperative role of chloride, bromide and peroxidasin in the assembly of the Col-IV^α121^ NC1 hexamer and formation of sulfilimine bonds in basement membranes. (A) Triple helical protomers of collagen IV are formed within the endoplasmic reticulum inside the cell, and are subsequently exported to the basement membrane. (B) Protomers oligomerize via their trimeric NC1domains forming Col-IV**^α121^** scaffolds that form the foundation of basement membranes. The assembly process is initiated by extracellular chloride pressure, which drives the oligomerization of individual trimeric protomers into hexameric structures (first arrow). Hexamers are subsequently stabilized and crosslinked into larger assemblies through the formation of sulfilimine bonds (second arrow) and the crosslinking of the N-terminal 7S domain (not shown). The formation of the sulfilimine bond is catalyzed by the enzyme peroxidasin along with cofactors bromide and hydrogen peroxide. The inset panels show molecular-level detail of the sulfilimine bond, specifically showing the covalent linkage between Hyl^211^ and Met^93^ residues located on adjacent subunits.

Unexpectedly, we recently found in a study of the Col-IV^α345^ scaffold, the major component of the kidney ultrafilter and the Goodpasture autoantigen, that sulfilimine bonds, independent of Cl- ions, also stabilized the hexamer quaternary structure. This finding revealed that the bonds covalently stabilized, not only the trimer-trimer interface of the NC1-hexamer, but also the side-by-side interactions of subunits within each trimer (Boudko et al., 2023; Pedchenko et al., 2019, 2021). Here, we sought to determine whether stabilization of hexamer quaternary structure by sulfilimine bonds also pertains to the Col4^α121^ scaffold that occurs ubiquitously across the animal kingdom, and whether the mechanism of bond formation, involving Br- and peroxidasin, are evolutionary conserved.

We found that sulfilimine bonds covalently reinforce the quaternary structure of the Col-IV**^α121^** hexamer, imposed by chloride conformational constraints. The bonds covalently fasten a clasp-motif across the trimer-trimer interface, interlocking the domain-swapping region of neighboring subunits, which reinforces the hexamer quaternary structure imposed by chloride conformational constraints. Collectively, our findings reveal that the structural reinforcement by sulfilimine bonds is a critical event in Col-IV scaffold assembly, enabling multicellularity, evolution and adaptation of metazoans, beginning with ancient cnidarians.

This paper is part of a series of publications on the chemistry and biology of collagen IV in the Journal of Biological Chemistry. Previously, we explored the essential role of chloride pressure in the assembly and stability of the collagen IV scaffold (Boudko et al., 2023) and uncovered the mechanism behind collagen IV assembly in *Drosophila* (Summers et al., 2023). We elucidated the evolutionary origin of a kidney-specific isoform of collagen IV, that enabled the compaction and function of the kidney filter (Pokidysheva et al., 2023). We determined the evolutionary origin and diversification of collagen IV genes and discovered that multiple cysteine residues in the collagenous domain of the kidney-specific isoform confers mechanical strength to kidney filter (P. S. Page-McCaw et al., 2025). The current study reports that sulfilimine bonds are a fundamental structural feature of Col-IV scaffolds that enabled metazoan evolution.

## Results

### Sulfilimine bonds, independent of Cl-, stabilize the quaternary structure of the mammalian Col4^α121^ hexamer

To determine whether sulfilimine bonds, independent of Cl-, stabilize the quaternary structure of Col4^α121^ NC1 hexamers, we investigated hexamers derived from two bovine tissues that are composed of two levels of sulfilimine-crosslinked dimer subunits. Col4^α121^ NC1 hexamer from bovine lens-capsule basement membrane (LBM) is predominately composed of monomer subunits, denoted as uncrosslinked-hexamer (Fig. 2A and 2B; (R. M. Vanacore et al., 2004)), whereas hexamer from bovine placental basement membrane (PBM) is composed predominately of dimers, denoted as crosslinked-hexamer (Fig. 2D and E; (Boudko et al., 2023; Pedchenko et al., 2019)). Hexamers were purified from both basement membranes, and their composition was assessed by SDS PAGE and size exclusion chromatography (SEC). In the presence of Cl-, the uncrosslinked LBM hexamer eluted as a single peak by SEC under nondenaturing conditions (Fig. 2B), but on Cl- removal, the hexamer dissociated predominately into monomers; about ten percent, composed of dimers, was resistant to dissociation (Fig.2C). In contrast, the crosslinked PBM hexamer was resistant to dissociation upon removal of Cl-, with a small amount, that composed of monomers, dissociated into monomers (Fig. 2F). These results indicate that sulfilimine bonds, independent of Cl-, stabilize the Col4^α121^ NC1 hexamer.

**Figure 2.**
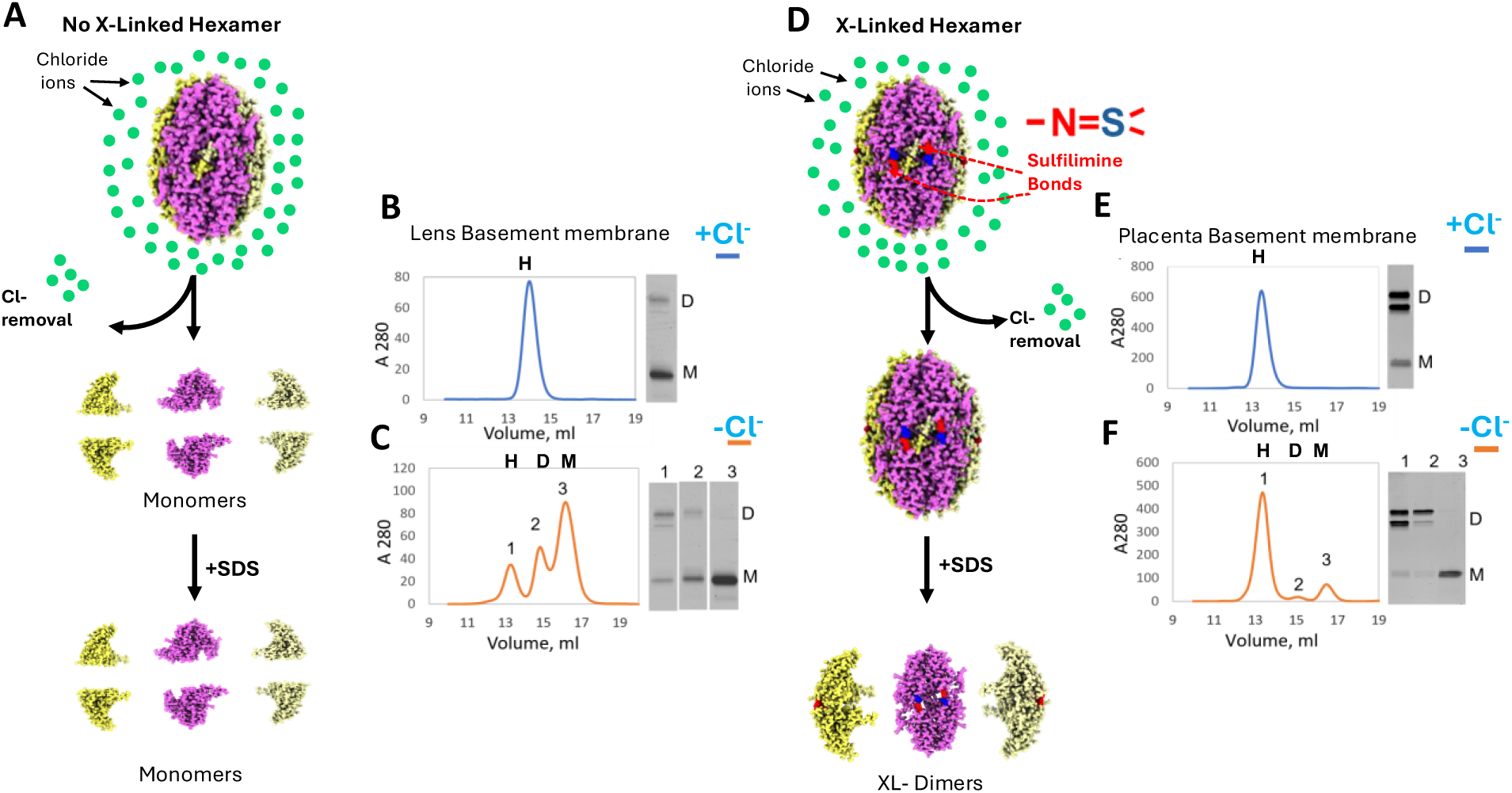
Chloride ions and sulfimine bonds, independently stabilize Col-IV ^α121^ NC1-hexamers in bovine lens and placenta basement membranes. Stabilization mechanisms of Col-IV ^α121^ hexamers derived from bovine lens basement membrane (LBM) versus placental basement membranes (PBM). (A) Schematic representation of LBM collagen IV hexamers, which contain a low level of sulfilimine crosslinks, relying primarily on chloride pressure for stability. Upon chloride removal, LBM hexamers undergo disassembly into monomeric subunits. (B) Size exclusion chromatography analysis showed LBM hexamers are stable in the presence of chloride, and are composed mainly of uncrosllinked monomer subunits as shown by SDS PAGE analysis. Hexamers are marked “H”, dimers “D” and monomers “M”. (C) Size exclusion chromatography and SDS-PAGE analysis of lens hexamers following chloride depletion, shows hexamer destabilization demonstrated by the increase in monomers and decrease in hexamers and dimers. (D) Schematic representation of placental collagen IV hexamers, which are stabilized by both chloride pressure and covalent sulfilimine bonds. These hexamers maintain structural integrity even after chloride removal due to sulfilimine bonds. (E) Size exclusion chromatography of placental hexamers in chloride-containing buffer with corresponding SDS-PAGE analysis shows that they are composed of crosslinked dimers. (F) Upon depletion of chloride, PBM hexamers remain structurally intact with only a minor amount of dissociation into monomers. SDS-PAGE confirms that the stable hexamer population consists primarily of crosslinked dimers, while the minor later-eluting peaks correspond to monomeric species.

To further investigate the role of sulfilimine bonds, independent of Cl-, on hexamer quaternary structure, we prepared Col4^α121^ NC1 hexamers from basement membrane in which the level of crosslinked-dimer subunits was experimentally altered by inhibition of peroxidasin activity. Previously, we demonstrated that mouse PFHR9 cells in culture produce collagen IV and that peroxidasin catalyzed sulfilimine bond formation. Treatment of cells with phloroglucinol (PHG), an inhibitor of peroxidasin (Bhave et al., 2012; McCall et al., 2014), altered crosslink formation. In the present study, hexamers were prepared from the PFHR9 matrix, grown in the presence or absence of PHG, and then were characterized by SEC and SDS PAGE. Control hexamers (crosslinked) are stable in the presence or absence of Cl- (Fig. 3A blue). In contrast, PHG-treated hexamers (uncrosslinked) are stable in the presence of Cl- but dissociate into monomers in the absence of Cl- (Fig. 3B red vs orange). We further explored the role of sulfilimine bonds by characterization of hexamers derived from the kidney and intestine of peroxidasin-knockout and wildtype mice. As expected, in the peroxidasin-KO animals, crosslinking is significantly reduced in these tissues, though not absent (Fig. 3C and D red vs blue). To determine whether this decreased crosslinking affects hexamer stability, we analyzed the NC1 fraction in the absence of Cl-. In both tissues, the NC1 hexamer derived from knockout animals ran predominantly as a monomer in SDS-PAGE gels in the absence of Cl-. Collectively, our results provide evidence (*in vitro and in vivo)* that sulfilimine bonds stabilize the hexamer quaternary structure in two mammalian species (*B. tauru*s and *M. musculus*), independent of Cl-, and corroborate our previous findings that peroxidasin catalyzes bond formation(Bhave et al., 2012) .

**Figure 3.**
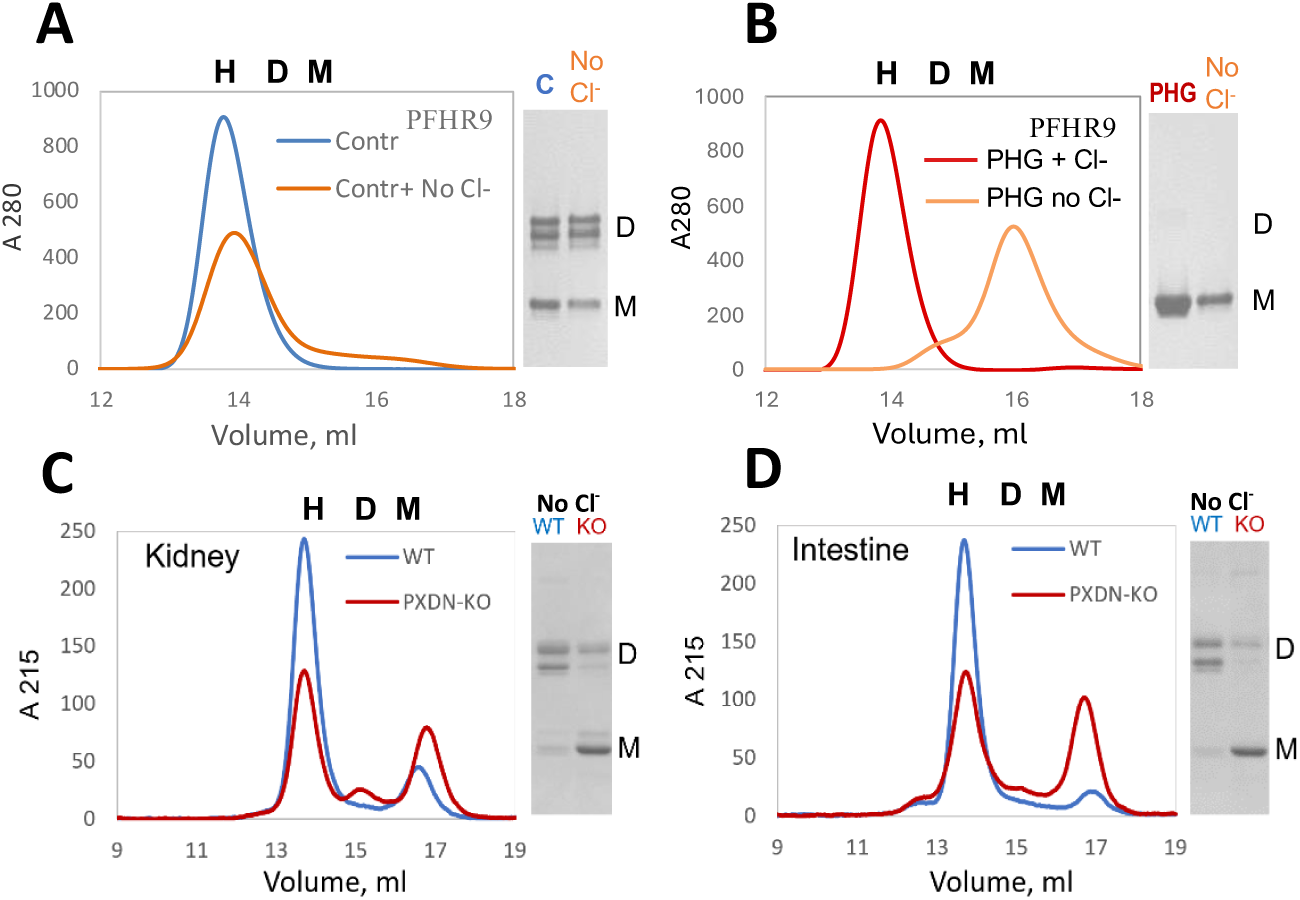
Sulfilimine bonds stabilize the quaternary structure of Col-IV ^α121^ NC1-hexamers against Cl- depletion matrix from PFHR9 cell culture and tissues of peroxidasin knock-out mice. (A) Size exclusion chromatography and SDS-PAGE analysis of NC1 hexamers purified from PFHR9 cell-derived extracellular matrix. NC1 domains maintain hexameric assembly both in the presence (blue trace) and absence (orange trace) of chloride, with corresponding SDS-PAGE analysis revealing predominantly crosslinked dimers under both conditions. Hexamers are marked “H”, dimers “D” and monomers “M”. (B) Analysis of NC1 domains purified from PFHR9 cells treated with the peroxidasin inhibitor PHG. Size exclusion chromatography shows hexameric NC1 assembly in the presence of chloride (red trace), but complete disassembly to monomers upon chloride removal (orange trace). SDS-PAGE confirms the absence of crosslinking in both samples, demonstrating the dependence on chloride for stability when sulfilimine bonds are inhibited. (C) Comparative analysis of NC1 domains purified from kidney tissue of wild-type mice (blue trace) versus peroxidasin knockout mice (red trace). Wild-type samples show predominantly hexameric NC1 with extensive crosslinked dimer formation, while peroxidasin knockout samples exhibit increased monomers and reduced crosslinking. (D) Analysis using intestinal tissue derived Col-IV NC1, where wild-type mice (blue trace) display stable hexameric NC1 with robust crosslinking, contrasted with peroxidasin knockout mice (red trace) showing elevated monomeric NC1 and diminished crosslinking capacity.

### Sulfilimine bonds, independent of Cl-, stabilize the quaternary structure of N. vectensis Col4^α121^ NC1 hexamer

In our previous studies, we discovered that the occurrence of the sulfilimine bond in Col4^α121^ hexamer, as well as the role of Cl- in hexamer stabilization, were conserved across animalia, beginning with cnidarians (Boudko et al., 2023; Fidler et al., 2014; P. S. Page-McCaw et al., 2025). Here, we sought to determine a) whether the quaternary-stabilization function of the bond, independent of Cl- *vide supr*a, and b) whether the mechanism of bond formation, involving peroxidasin and Br- (Bhave et al., 2012; McCall et al., 2014), pertain to basal cnidarians. To this end, we investigated *Nematostella* vectensis, a cnidarian that represents an early diverging metazoan, and they are among the simplest animals with a basement membrane (Tucker et al., 2011; Tucker & Adams, 2014). Recent work has revealed that they possess a complex and dynamic matrisome, offering a glimpse into the key evolutionary events that produced the BM (Bergheim et al., 2025). They also are a model organism with increasing number of established techniques that make them an important organism in which to study the function, structure and assembly of basement membranes, and particular collagen IV. (Bergheim et al., 2025; Carvalho et al., 2025; Layden et al., 2016; Paix et al., 2023; Röttinger, 2021).

We first characterized the structure of Col4^α121^ NC1 hexamer from *N.vectensis* and analyzed the NC1 oligomeric state by SEC. We excised the hexamer from the whole animal by collagenase digestion and then purified the hexamer by gel exclusion chromatography (Fig. 4A, red arrow). We noted that the hexamer subunits from *Nematostella* had similar electrophoretic mobility as those derived from bovine lens capsule BM (Fig. 4B). We confirmed the identity of the protein bands observed with Coomassie stain by two independent means. First, we demonstrated that these bands cross-reacted with anti-Col IV NC1 domain immune serum in a Western blot (Fig. 4B right). Second, we analyzed tryptic peptides from each band for peptide sequence by mass spectroscopy, and found peptides corresponding to both an α1 and an α2 Collagen IV NC1 sequence from *Nematostella* (Fig. 4C and D). Together these results demonstrate that *Nematostella* contains a collagen IV scaffold composed of α1 and α2 chains with an NC1 domain that migrates on denaturing SDS-PAGE as monomers and dimers similar to those found in mammals.

**Figure 4.**
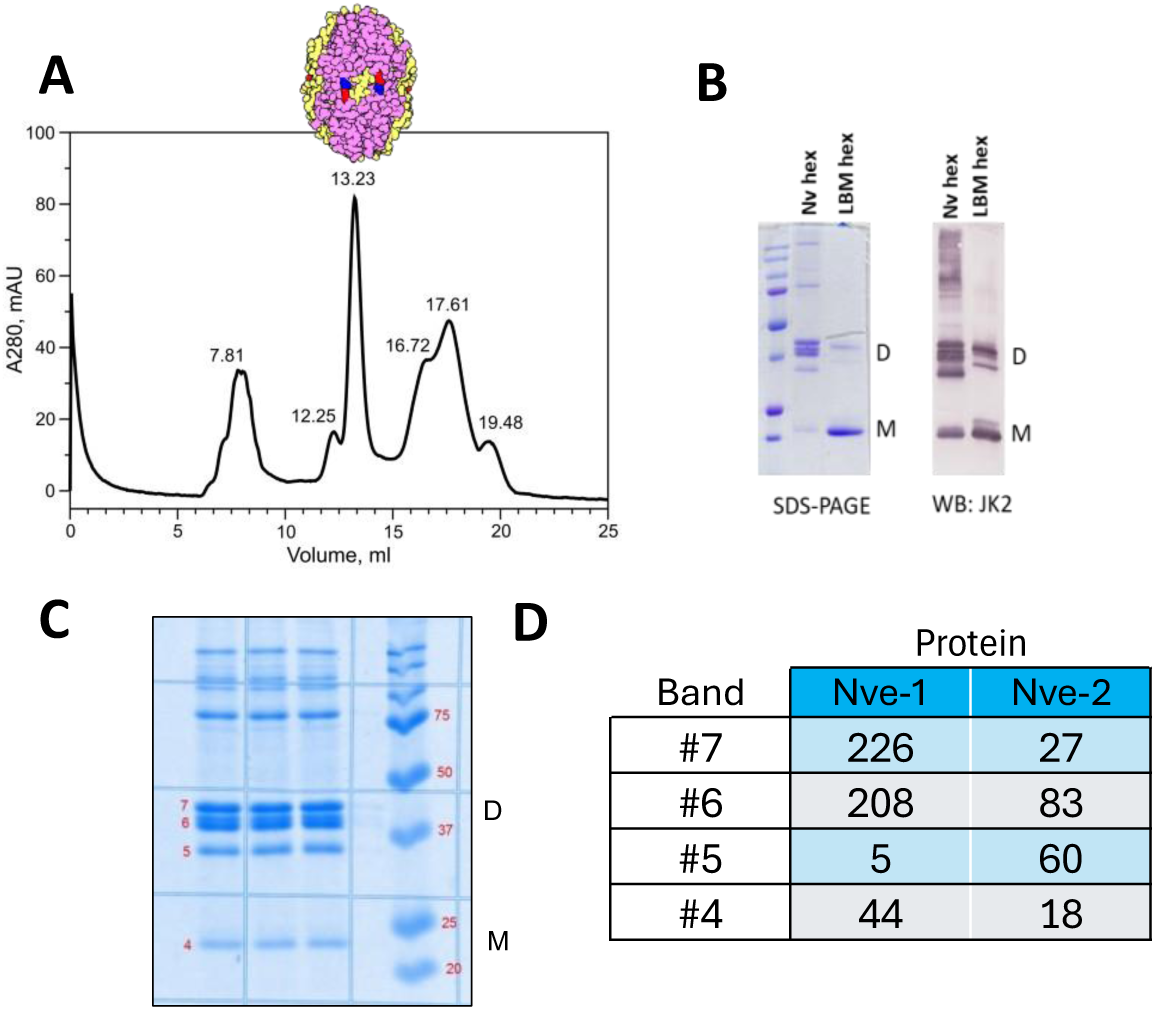
Purification and characterization of Col-IV ^α121^ NC1-hexamer from N. vectensis. Evolutionary conservation of collagen IV structure and assembly mechanisms. (A) Gel-filtration FPLC purification profile showing the separation of NC1 hexamers from collagenase-digested *N. vectensis* tissue, with the red arrow indicating the FPLC peak fraction corresponding to the NC1 hexamer. (B) SDS-PAGE and Western blot analysis confirming the successful isolation of Col-IV NC1 hexamers from *N. vectensis* and demonstrating their subunit composition, including both crosslinked dimeric subunits “D” and uncrosslinked monomeric subunits “M”. Purified NC1 hexamer from bovine lens basement membrane (LBM) serves as a positive control for comparison. (C and D) Confirmation of the expression of two distinct alpha chain types (α1 and α2) in *N. vectensis* Col-IV NC1 domains at the translational level. Mass spectrometry analysis of isolated *N. vectensis* NC1 hexamers shows the presence of two α1- and α2-like NC1 domain sequences at the protein level, with sequence coverage of 52-58% for both chain types.

We next characterized the primary and hexamer structure of the Col4 NC1 domains of *Nematostella*. First, we performed a sequence alignment of the Col4 NC1 domains of *Nematostella* with humans and several model organisms to determine the level of conservation of the hydroxylysine-211 and methionine-93 (Fig. 5A) residues required for formation of the sulfilimine bond. We found that these residues were absolutely conserved across all of the aligned species (Fig 5B). Further, we performed predictions of the structure of NC1 from *Nematostella* using AlphaFold 3 (Fig. 5C) and compared this prediction to the crystal structure of the human NC1 (Fig. 5D, PDBID 1LI1) (Than et al., 2002). Overall, the structures were similar, and crucially, the conserved lysine 211 and methionine 93 were predicted to be aligned as if they were crosslinked despite the prediction being conducted in the absence of any crosslinking information.

**Figure 5.**
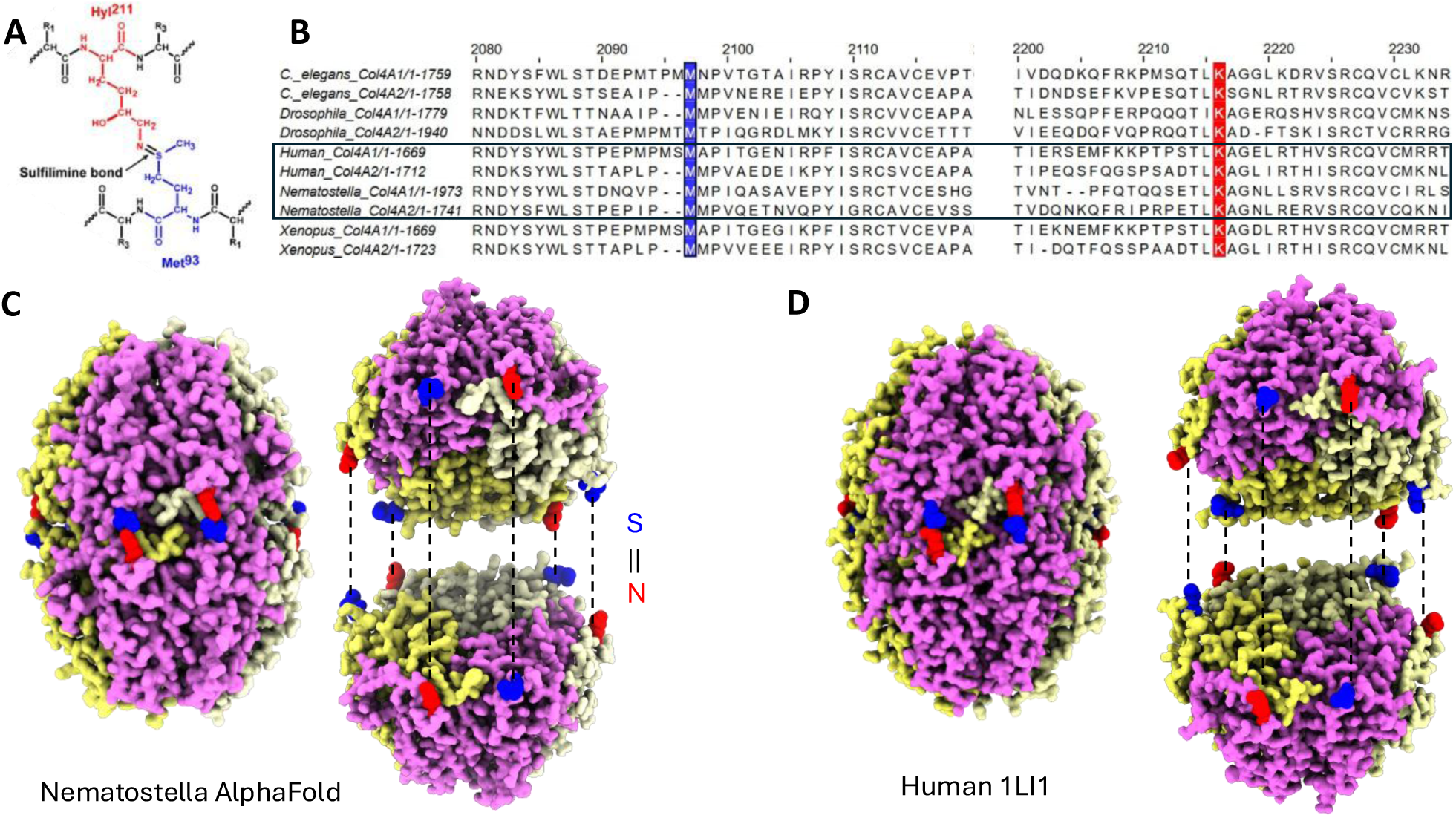
The residues required for sulfilimine bond formation are conserved from *Nematostella* to humans, and are predicted to be aligned with each other based on alpha fold 3. (A) Molecular detail of the sulfilimine bond showing the covalent linkage between Hly ^211^ (red) and Met^93^ (blue), with the characteristic double bond between nitrogen and sulfur atoms labeled. (B) Multiple sequence alignment of Col-IV α1 and α2 chains from *C. elegans*, *D. melanogaster*, *H. sapiens*, *N. vectensis*, and X*. laevis*. The sequences from human and *N. vectensis* are indicated by a box, and the critical methionine (blue) and lysine (red) residues required for sulfilimine bond formation are marked. These residues show conservation across all examined species. (C) AlphaFold 3 structural prediction of *N. vectensis* Col-IV ^α121^ NC1 hexamer, demonstrating remarkable structural similarity to the experimentally determined human structure. Notably, the lysine and methionine residues essential for sulfilimine bond formation are predicted to be positioned in close proximity to each other in a configuration compatible with crosslink formation. (D) Crystal structure of human Col-IV NC1 hexamer (PDB: 1LI1) shown for comparison, highlighting the structural conservation between cnidarian and mammalian Col-IV assemblies. More details from the AlphaFold3 prediction are in Fig. S1.

Next, we investigated the conservation of the 3D structure of peroxidasin, the enzyme that catalyzes formation of the sulfilimine bond (Bhave et al., 2012; Ero-Tolliver et al., 2015). Because there are no high-resolution structures of peroxidasin, AlphaFold 3 was used to predict the structures of the catalytic domains of both human and *Nematostella* peroxidasin to investigate conservation. AlphaFold 3 predictions of the human (Fig. 6A, B, and C) protein were very similar to the predicted structure of the *Nematostella* protein (Fig. 6D, E, and F). Interestingly, in both predictions there are three pairs of cystine residues that are highly conserved and are predicted to be placed in proximity to form disulfide bonds that would stabilize the trimeric enzyme (Fig. 6B and E, purple residues with yellow side chains). Additionally, both enzymes are predicted to bind to heme at highly conserved binding pockets, and the key residues for trimerization and heme binding are highly conserved (Fig 6. B, C, E, and F, purple residues).

**Figure 6.**
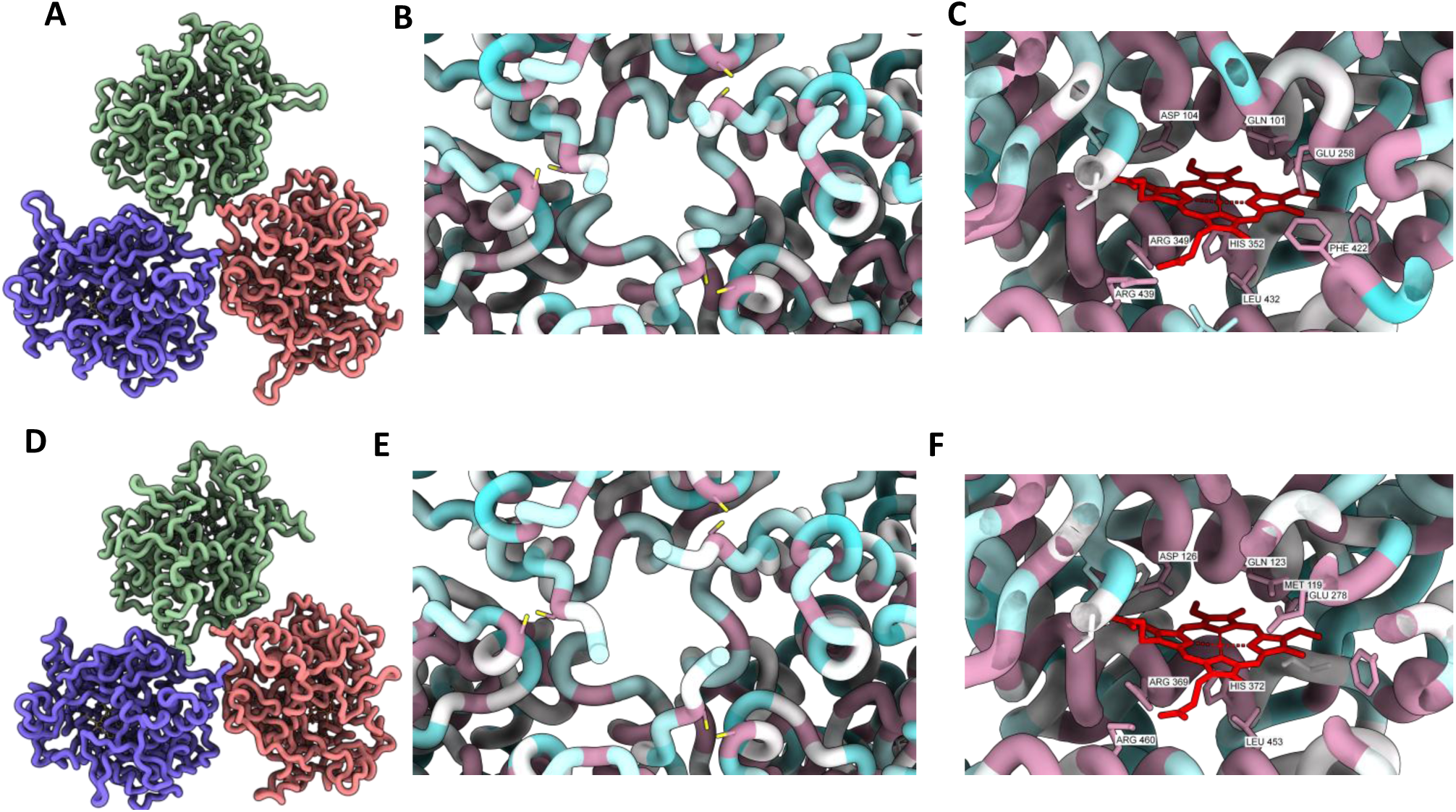
The core features of peroxidasin are conserved from *Nematostella* to humans, including the trimerization domain and the heme binding site. AlphaFold structural predictions of human (A) and N. vectensis (D) peroxidasin trimers, with individual chains shown in different colors. Detailed views of the trimerization interface regions in human (B) and N. vectensis (E) peroxidasin predictions, with residues colored according to evolutionary conservation based on Clustal Omega alignment analysis of peroxidasin sequences from *H. sapiens*, *X. laevis*, *D. melanogaster*, *C. elegans*, and *N. vectensis*. Highly conserved residues are displayed in magenta, while more variable residues are shown in cyan. The trimerization interface of both human and *Nematostella* contain 3 pairs of conserved cysteine residues. Analysis of heme binding sites in human (C) and *N. vectensis* (F) peroxidasin AlphaFold predictions, with residue coloring based on the same conservation analysis. The heme coordination environment is maintained by several highly conserved residues (magenta). More details of the AlphaFold prediction and alignment for color scheme are found in Fig. S2 and S3.

We next determined whether Br- is required for sulfilimine bond formation and stabilization of Col4^α121^ NC1 hexamers in N*ematostella*. Because Br- is absolutely required for sulfilimine bond formation as a cofactor for peroxidasin in mammals and drosophila, (Bhave et al., 2012; Colon & Bhave, 2016; Ero-Tolliver et al., 2015; McCall et al., 2014), we grew and maintained *Nematostella* in Br-free media, prepared NC1 hexamers, and then determined their level of degree of crosslinked-dimer subunits by SDS-PAGE (Fig. 7A), as a measure of bond formation. Under these conditions, the amount of sulfilimine bond formation is greatly reduced, though not absent (Fig. 7A, compare lane 1 to 3, quantified in 7B). The remnant sulfilimine crosslink may have had several sources. First, it may represent preexisting crosslinked BM present in these animals. Collagen IV is known to be a long-lived molecule and has been shown to have a long half-life in vivo. Here, the animals grew slowly, suggesting that much of the BM at harvest was present at the time when the animal was moved to Br- free conditions. Second, while the animals were maintained in the absence of Br- in the media, their food source may have contained some residual Br- sufficient to prevent a complete block to sulfilimine bond formation, despite efforts being made to avoid this. Finally, a low level of crosslinking may occur by an alternative mechanism as hinted by the presence of sulfilimine bonds observed in Pxdn KO mice (Fig. 3C and D). Nonetheless, crosslinking is greatly reduced in *Nematostella* grown under the Br- free condition, revealing a critical role of Br- as a cofactor of the peroxidasin-catalyzed formation of the sulfilimine bond in a basal animal.

**Figure 7.**
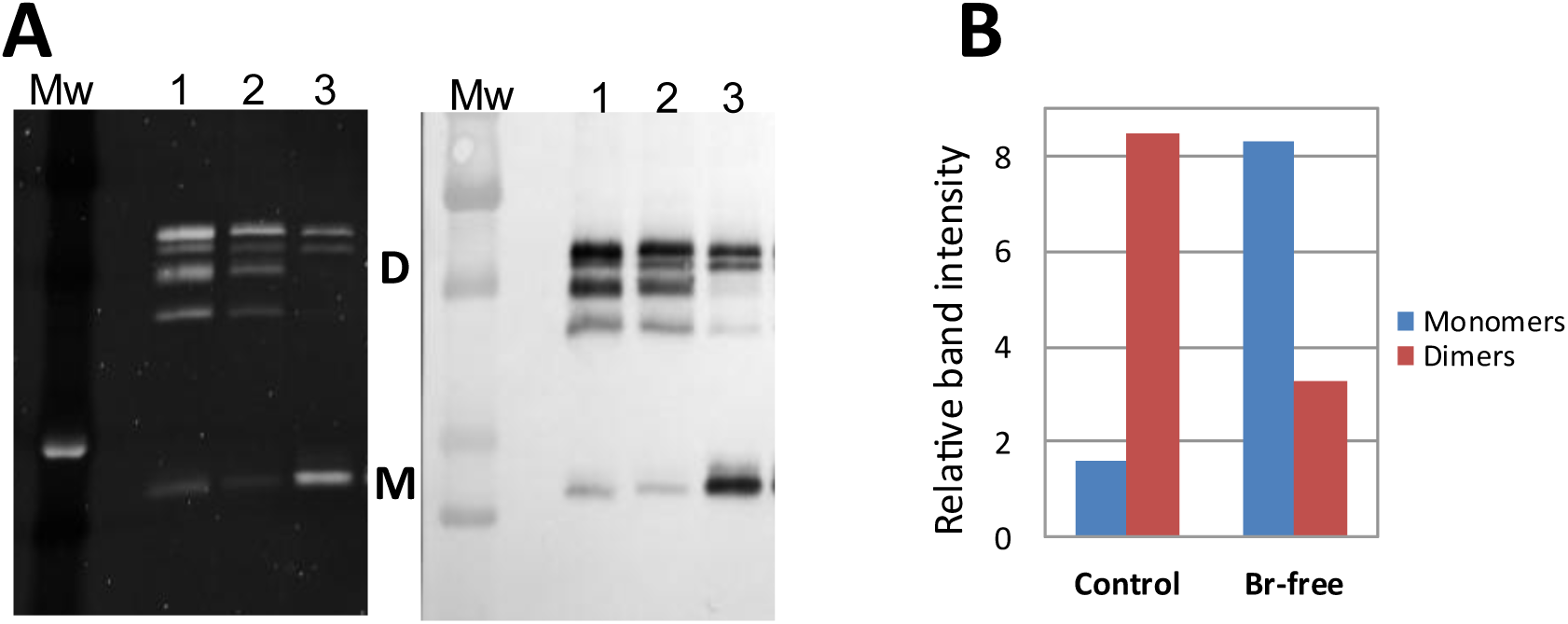
Bromide is a critical ion for the formation of sulfilimine bonds in Col-IV^α121^ scaffolds of *Nematostella*. (A) SDS-PAGE and Western blot analysis of isolated NC1 comparing crosslinking patterns between control and bromide-depleted conditions. Lane 1 shows control purified N. vectensis NC1, lane 2 displays NC1 from animals cultured in standard control media, and lane 3 presents NC1 from animals cultured in bromide-free medium. Protein samples were analyzed by SDS-PAGE with SYPRO Ruby staining (first panel) and by Western blot (second panel) using the Col-IV-specific JK2 antibody. Bromide depletion results in a significant increase in monomeric NC1 species “M” compared to control conditions, with a reduction in crosslinked NC1 dimers “D”, indicating impaired sulfilimine bond formation in the absence of bromide. (B) Quantitative analysis of NC1 monomer and dimer band intensities performed by gel densitometry.

We next determined whether Br- plays a direct role in *Nematostella* hexamer stability, given that Cl- directly stabilizes hexamer quaternary structure in Nematostella and across the animal kingdom (Boudko et al., 2023; Cummings et al., 2016; Pedchenko et al., 2019). We recently developed a simple *in vitro* method of SDS-PAGE to determine the impact of Cl- on hexamer stability (Pedchenko et al., 2021) With this method, Nematostella NC1- hexamer was analyzed first by its migration on traditional SDS PAGE (Fig. 8A, No Cl^-^ gel) then compared with its migration on a gel containing 150mM NaCl in both the gel and running buffer (Fig. 8B, +Cl^-^ gel). In the no Cl- gel (Figs. 8A), hexamers dissociate into dimers and monomers, whether the sample is at room temperature (RT) or boiled (B) prior to running the gel. In hexamers prepared under Br-free conditions, the amount of dimers is greatly diminished, whereas monomers is greatly increased (Fig. 8A), confirming a critical role of Br- in bond formation as described in Fig.7B. However, in the + Cl- gel and running buffer, hexamers prepared under control and Br-free conditions both migrate as intact hexamers at RT but dissociate by boiling into its dimer and monomer subunits (Fig. 8B), as depicted in Fig. 8C. These results reveal that Br-, in contrast to Cl-, does not directly stabilize hexamer quaternary structure.

**Figure 8.**
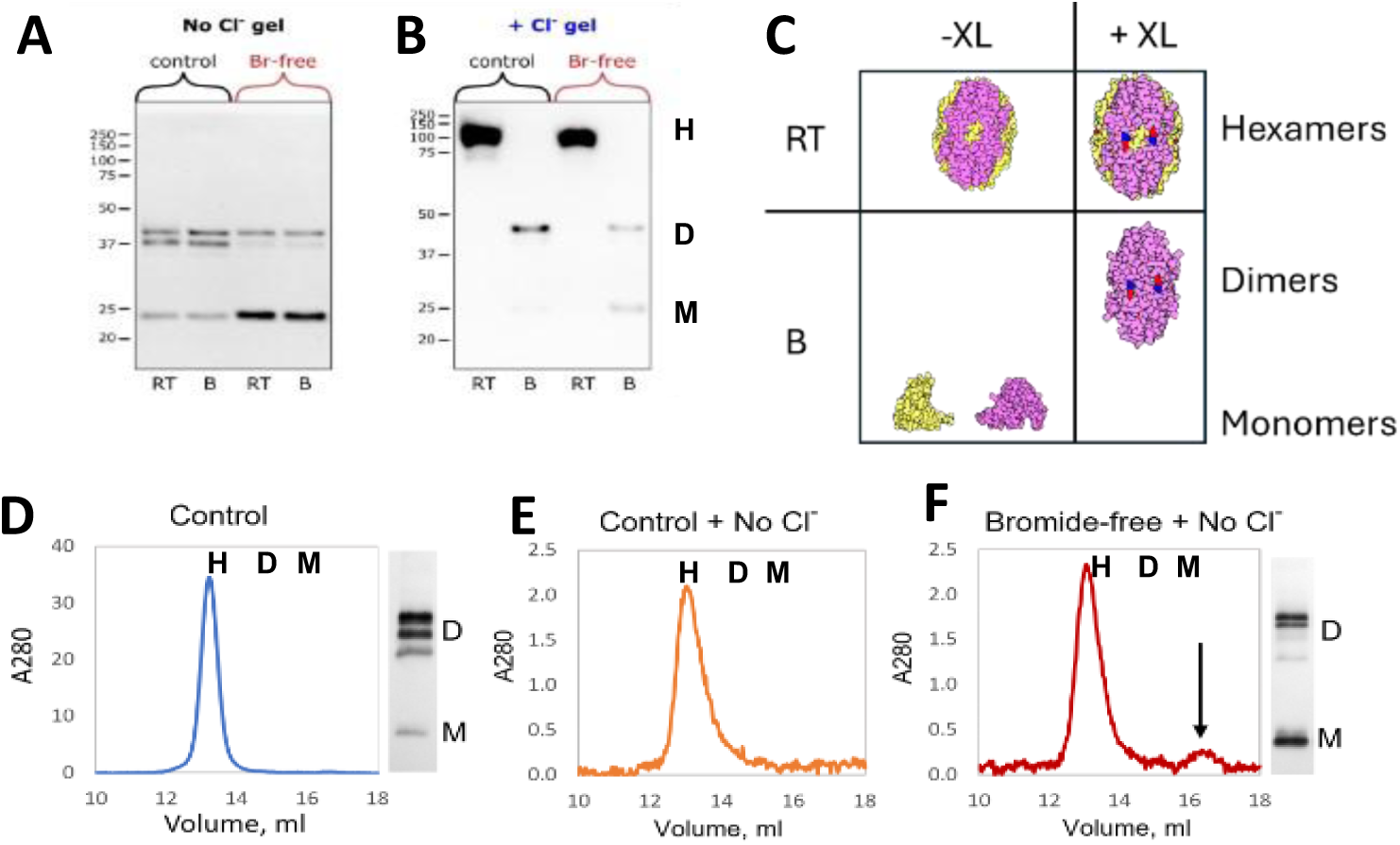
Sulfilimine bonds, independent of Cl-, stabilize the hexamer quaternary structure in *Nematostella*, as was found for mammals. (A) SDS-PAGE analysis of *N. vectensis* NC1 samples under chloride-free conditions, comparing control animals (normal growth conditions) with animals cultured in bromide-free medium. Samples analyzed at room temperature “RT” and after boiling “B” reveal that bromide depletion reduces crosslinked dimers “D” and increases monomers “M”, indicating impaired sulfilimine bond formation. (B) Parallel analysis under chloride-containing gel conditions shows that both control and bromide-depleted samples maintain hexameric “H” assembly at room temperature. However, following boiling, control samples show predominantly crosslinked dimers, while bromide-free samples exhibit a mixture of dimers and monomers, confirming reduced crosslinking stability. (C) Schematic representation illustrates the expected results from Chloride gel analysis of crosslinked and uncross linked samples under RT and boiled conditions. (D) Size exclusion chromatography of control *N. vectensis* NC1 in chloride-containing buffer shows stable hexamers, with SDS-PAGE analysis of the peak fraction revealing predominantly crosslinked dimer composition of the hexamers. (E) Analysis of control NC1 sample under chloride-free conditions shows maintained hexameric assembly, indicating that sulfilimine crosslinks alone are sufficient to stabilize hexamer structure in the absence of chloride pressure. (F) Size exclusion analysis of bromide-depleted N. vectensis hexamer under chloride-free conditions shows stabilized hexamers with a small amount that dissociated into monomers. SDS-PAGE analysis showed decreased dimer prevalence and increase in monomers, demonstrating that the residual amount of bond formation was sufficient to confer resistance to hexamer dissociation when chloride is removed.

We next determined whether sulfilimine bonds, independent of Cl-, stabilize the quaternary structure of the Col4^α121^ NC1 hexamer in *Nematostella.* Using SEC as described in Figs. 2 and 3 for mammals, we found that in *Nematostella* the hexamers are predominately crosslinked and are stable in the presence (Fig. 8D) and absence of Cl-(Fig. 8E). Likewise, hexamers assembled under Br-free conditions are stable upon removal of Cl-, despite reduced levels of dimers, and only a small amount dissociate into monomers on removal of Cl- (Fig. 8F, arrow). This finding is analogous with that for the bovine lens capsule hexamer in which a small amount of dimer confers stability to hexamer upon removal of Cl- (Fig. 2). Collectively, these results indicate that sulfilimine bonds, independent of Cl-, stabilize the hexamer quaternary structure in *Nematostella*, as was found for mammals.

### Sulfilimine bonds and mechanism of formation are conserved in collagen IV across metazoan

Evidence for phylogenetic conservation of the sulfilimine bond and mechanism of formation in Col-IV scaffolds is summarized in Fig. 9A. Dimer subunits of NC1-hexamers, crosslinked by sulfilimine bonds (Fig.9B), occurs in species across Animalia. Mass spectrometric analyses verified the crosslinks in key species (Fig. 9A (Fidler et al., 2014)). Furthermore, the mechanism of bond formation is conserved, as evidenced by a) phylogenic analysis of peroxidasin (Colon & Bhave, 2016; Fidler et al., 2014) b) the amino acid residues required for bond formation (Fig. S4), and c) the requirement for Br- in key species (Fig. 9A). Collectively, this evidence illuminates an essential role of the sulflimine bond in the evolution of multi-cellular animals, and particularly the essentiality of Br- in the formation of the sulfilimine bond.

**Figure 9.**
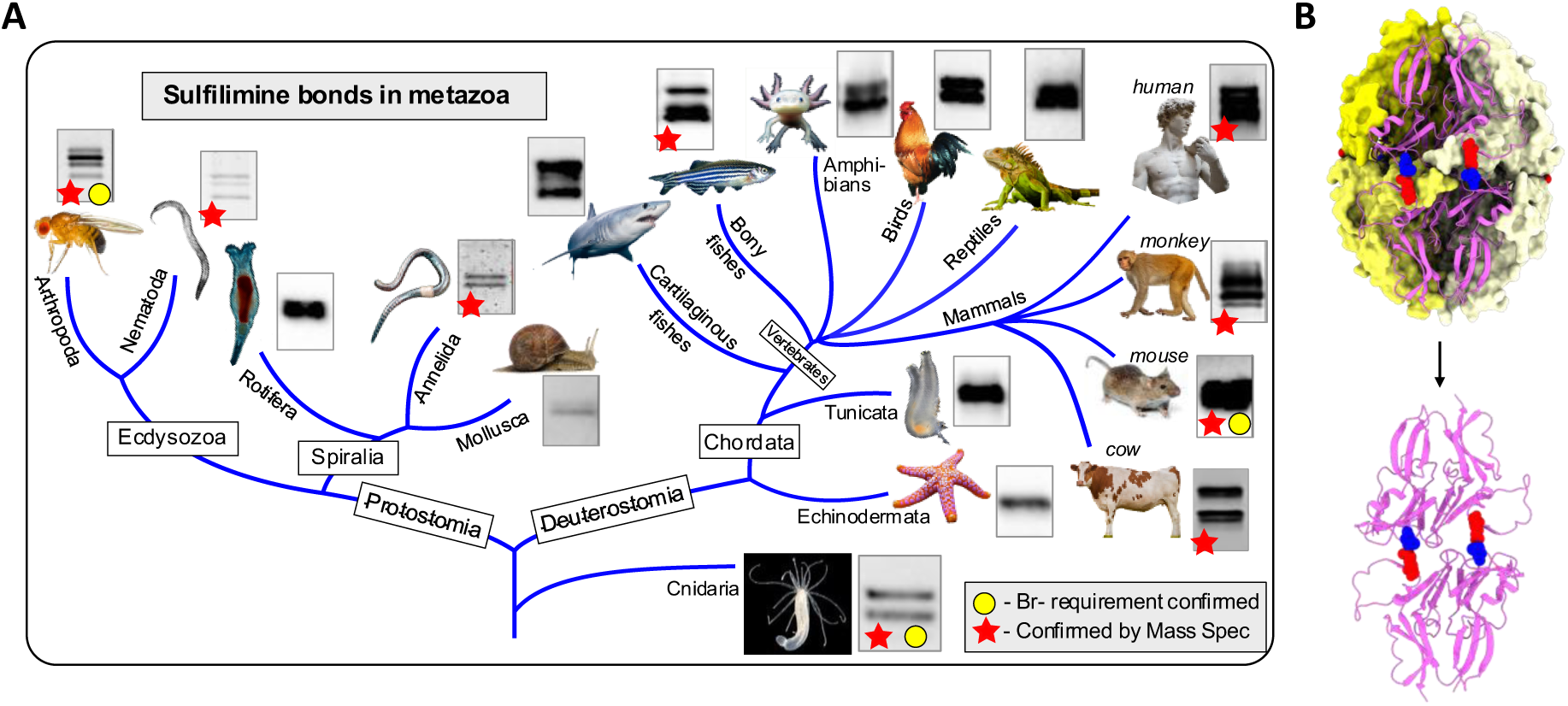
Bromine-mediated sulfilimine bonds are conserved across metazoan evolution, beginning with cnidarians. (A) Phylogenetic tree summarizes the experimental evidence for the evolutionary conservation of the occurrence of the sulfilimine bond across key metazoans. Red stars denote species in which the bond was confirmed by mass spectrometry. Yellow circles denote species in which Br- impacted bond formation. Each species node includes a representative SDS-PAGE gel section demonstrating the presence of crosslinked dimers, which serves as biochemical evidence for sulfilimine bond formation. The summary reveals that sulfilimine bond formation spans across the animal kingdom, beginning with the basal cnidarians. (B) Structural comparison showing a crosslinked NC1 hexamer and a crosslinked dimer subunit to illustrate the relationship between dimers under denaturing conditions and the presence of covalent sulfilimine bonds.

### Sulfilimine bonds function as a reinforcement of hexamer quaternary structure by interlocking the domain-swapping region of neighboring subunits

We next analyzed the crystal structure of the Col-IV**^α121^** hexamer to gain insight into the mechanism of how sulfilimine bonds increases stability of the hexamer quaternary structure. In assembly, hexamer formation and stability of the quaternary structure is driven by the extracellular chloride concentration - “chloride pressure”, which imposes conformational constraints within NC1-trimers and across the trimer-trimer interface (Fig. 1 first arrow, Figs. 5C and D) (Boudko et al., 2023; Cummings et al., 2016; Pedchenko et al., 2019). In a subsequent step, hexamer structure is further stabilized by the formation of six sulfilimine bonds. The bonds form covalent crosslinks at the hexamer interface that weld together trimeric NC1-domains of adjoining protomers (Fig. 1, 5C & D) (Boudko et al., 2021; McCall et al., 2014; R. Vanacore et al., 2009) Furthermore, the crosslinks stabilize the side-by-side interactions of neighboring subunits within each trimer, as described *vide supra* (Boudko et al., 2023; Pedchenko et al., 2019, 2021). Collectively, the covalent sulfilimine crosslinks increase the avidity of binding interactions across all six subunits, stabilizing the hexamer quaternary structure (Fig. 10). An understanding of the structural basis for how sullfilimine bonds increase avidity of interactions that stabilize the hexamer quaternary structure hinges on gaining insights about the secondary and tertiary structure of trimer subunits and their connection across the hexamer interface.

**Figure 10.**
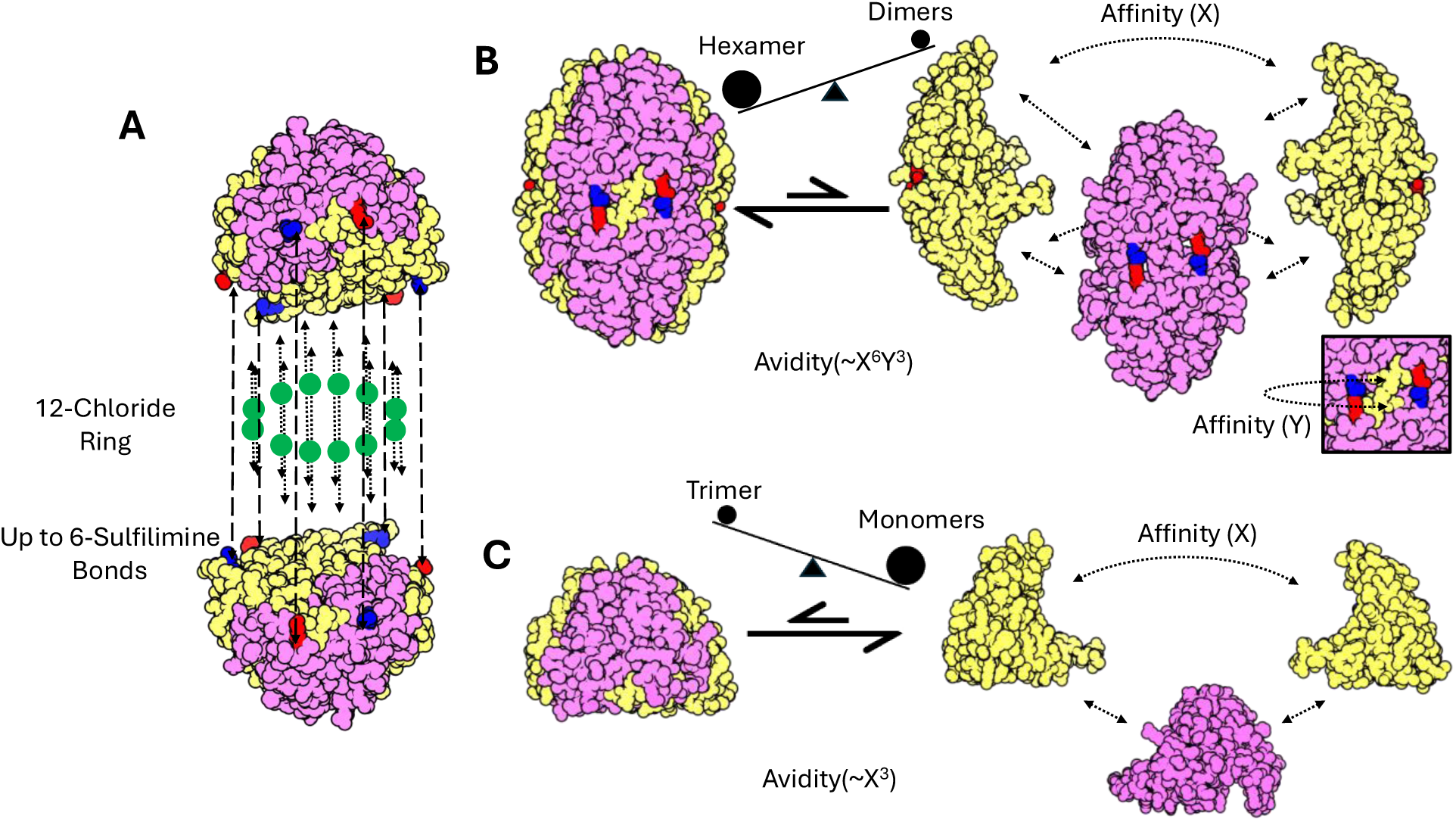
Affinity and avidity dynamics of NC1 trimers and hexamers. (A) The Col-IV**^α121^** NC1 hexamer is stabilized by two distinct chemical mechanisms, the 12 chlorides ions that form a ring structure between the 2 trimers, and the 6 sulfilimine bonds that form across the trimer-trimer interface. (B) In the crosslinked hexamer there are a number of different interactions across the six subunits, and the affinity of each of these interactions contribute to the overall avidity of the complex. The presence of the sulfilimine bond results in enhanced stability of the complex. This results in the hexamer being the preferred state (C) The trimer does not benefit from the same enhanced interactions, and thus prefers the monomeric state.

The overall structure of the trimer subunit consists of two rings, an upper and lower ring, each composed of a series of β-sheets. The lower ring of the NC1-trimer (marked in Fig. 11A and shown stylized in Fig. 11B) can be viewed as a β-propeller structure (Than et al., 2002) comprised of 6 blades, which resembles the β-pinwheel structure of the DNA Gyrase family (Corbett et al., 2004; Kopec & Lupas, 2013). Each blade is a 6-stranded β-sheet, composed of 4-strands from one subunit and a 2-stranded β-hairpin structure, stabilized by a disulfide bond, from the neighboring subunit or neighboring C4 domain (Fig. 11B, blades are numbered i to vi). The β-sheet-motif functions as a domain-swapping mechanism that connects neighboring subunits within trimers (Casino et al., 2018; Sundaramoorthy et al., 2002; Than et al., 2002). The sulfilimine bonds are located on the outer strand of each of the six blades. This placement constrains the lateral association of the neighboring subunits in two ways. First, the presence of the bond anchors the outer loop of blade ii which sterically constrains the domain swap from the neighboring subunit (light yellow insert into blade ii Fig. 11C and D). Second, the covalent stabilization of the outer beta strand (Fig 11E) means that the entire beta sheet of blade ii is now stabilized enhancing the interaction between the two neighboring subunits.

**Figure 11.**
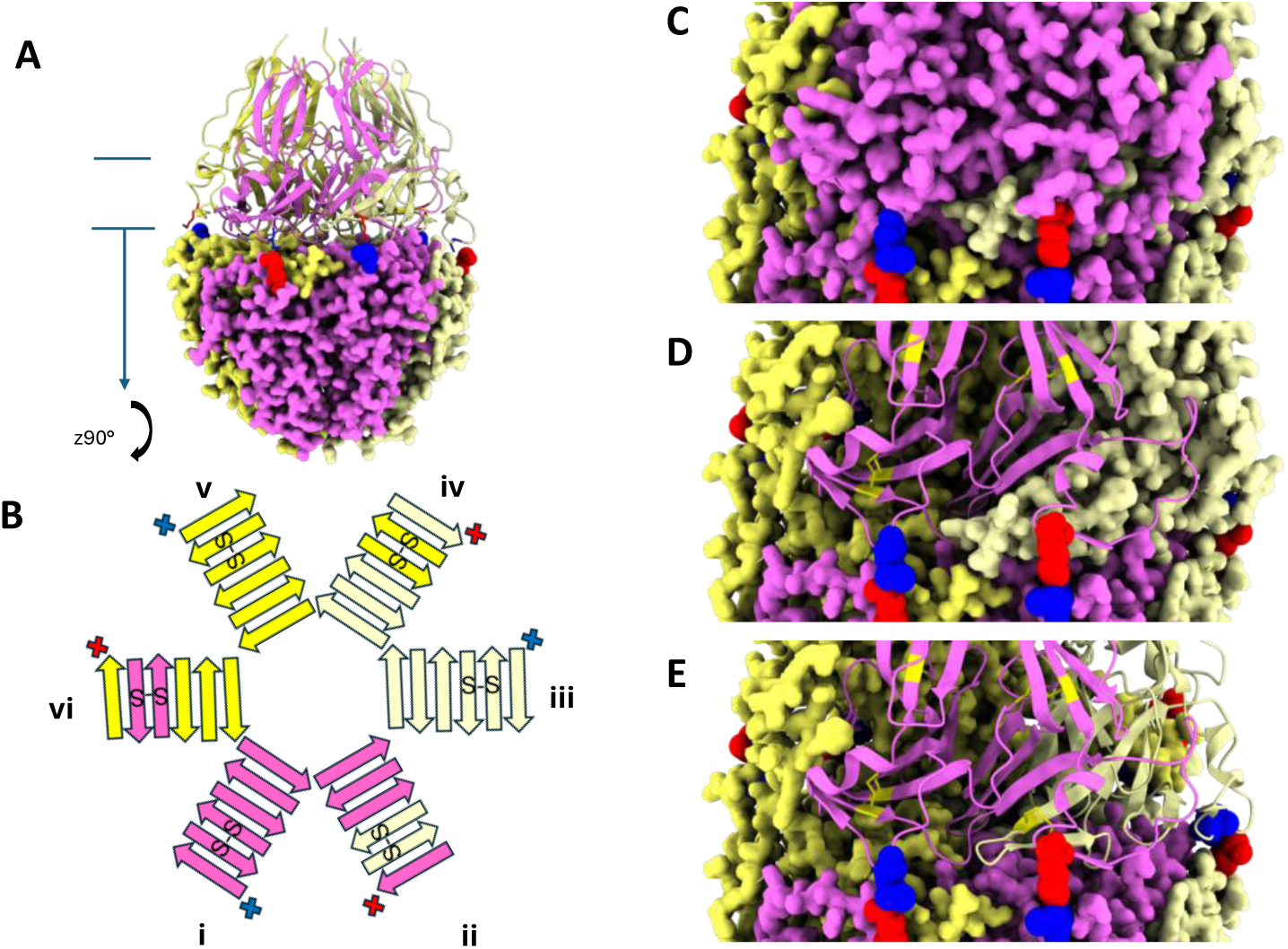
The sulfilimine bond, positioned at the terminus of a β-sheet structure, maximize the stabilization of the secondary structural elements of Col-IV^α121^ NC1 hexamers. (A) Structural representation of the human Col-IV**^α121^** NC1 hexamer with the lower protomer displayed in space-filling representation and the upper protomer in ribbon representation. The sulfilimine bond is highlighted with key lysine residues shown in red and methionine residues in blue, demonstrating the positioning of the crosslink at the trimer-trimer interface. (B) Stylized schematic representation of the β-pinwheel structure observed in the lower portion of the NC1 trimer including the location of disulfide bonds present in the β-hairpin of the domain swap. This schematic illustrates how the sulfilimine bonds work in concert with disulfide bonds to create a robust interlocking system that maintains hexamer stability. The six blades of the structure are numbered “i” to “vi”. Disulfides are represented as “S-S”. Sulfilimine residues are shown in red and blue. (C) An alternative view of the hexamer showing the placement of the sulfilimine bond and highlighting the location of the bond relative to the domain swap region from adjacent subunits (D) The same view as shown above, but with one of the chains shown as a ribbon. This highlights how the domain swap region from the adjacent subunit is sterically constrained by the outer loop stabilized by the sulflimine bond. Disulfide bonds are shown in yellow (E) The same view shown above, but with two subunits now shown as ribbons. This highlights how the constrained domain swap is a part of the β sheet. This sheet concludes with the outermost β strand located adjacent to the sulfilimine bond.

Within the hexamer, sulfilimine bonds covalently interlock the domain-swapping region of juxtaposed trimers, reinforcing the trimer-trimer interactions as well as the lateral associations of neighboring subunits (Fig. 12). These functions are coordinated in a region we define as the clasp-motif. The clasp-motif is formed by the outer portions of two blades of the β-propeller, noted as i and ii and characterized by two distinct side-by-side conformations. The clasp motif surrounds the β-hairpin domain swaps from blade iii inserted into the clasp. Sulfilimine bonds covalently connect i and ii of the top trimer, across the trimer-trimer interface, to ii* and i* respectively of the juxtaposed bottom trimer (Fig. 12B). Blade i and ii* bind the inserted β-hairpin iii’ from the neighboring subunit, and blades ii and i’ bind to β-hairpin iii (Fig. 12B). A sulfilimine bond covers the top of each β-hairpin structure which prevents trimer-trimer dissociation and sterically hinders lateral dissociation of neighboring subunits (Figs. 12B and 11C and 11D). In the absence of the bond, experimental removal of Cl- causes hexamer dissociation; however, when the clasp is locked by the sulfilimine bond, the hexamer quaternary structure is preserved (Figs. 2 and 12C). Taken together, the sulflilimine bonds covalently interlock the trimer-trimer interface, reinforce the β-pinwheel structure of trimers, and reinforce the lateral association of neighboring subunits within the trimer, which increases the avidity of binding interactions across all six hexamer subunits. In summary, the six sulfilimine bonds, located around the circumference, function as a covalent reinforcement of the hexamer quaternary structure.

**Figure 12.**
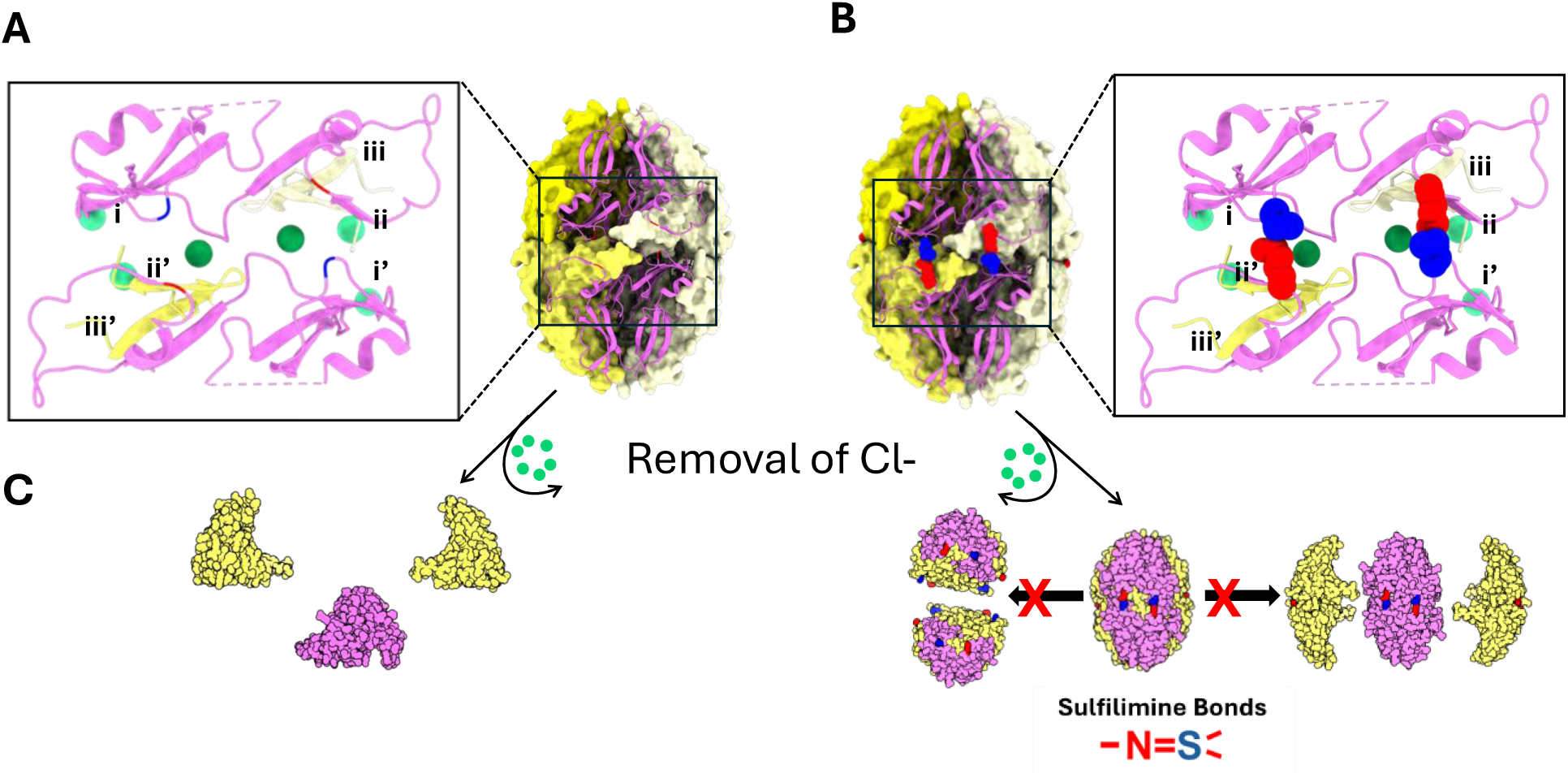
Six sulflilimine bonds covalently interlock the domain-swapping regions via a clasp-motif that reinforces the quaternary structure of the Col-IV^α121^ NC1-hexamer. (A) The “clasp motif” is positioned at the interface between two NC1 trimers, whose interactions are stabilized by the presence of chloride. Each clasp contains parts from 4 of the 3 alpha chains that make up the hexamer, including the sulfilimine residues from blades i and ii (denoted in Fig 11.) from one of the two primary alpha chains, i’ and ii’ from the other primary alpha chain, and the domain swap beta hairpins from blade iii and iii’ of the other two neighboring alpha chains. (B) The sulfilimine bond forms across the two primary alpha chains that compose the motif. This covalently links these two alpha chains, as well as providing significant stabilization of the neighboring domains swaps by providing conformational constraints on the outer beta strand of each beta pinwheel blade. (C) When chloride is removed from the un crosslinked hexamer, the complex collapses into monomers, however when the sulfilimine bond is present, the clasp motif holds the hexamer together

## Discussion

In this study, we gained new insights about the essentiality, molecular assembly and function of the unique sulfilimine bond in collagen IV scaffolds (Fidler et al., 2014; R. Vanacore et al., 2009). The bond, a sulfur-nitrogen bond (-S=N-), covalently holds together the Col-IV**^α121^**, Col-IV**^α556—α121^**, and Col-IV**^α345^** scaffolds (Hudson et al., 2003; Khoshnoodi et al., 2008; R. M. Vanacore et al., 2004). The Col-IV**^α121^** scaffold, the most well-characterized scaffold, is ubiquitously expressed across all animal phyla, imparting functionality to basement membranes of multicellular epithelial tissues (P. S. Page-McCaw et al., 2025). This scaffold provides mechanical strength, serves as a ligand for integrins and other cell-surface receptors, and interacting with growth factors such as BMPs to establish signaling gradients (Wang et al., 2008). Col-IV**^α121^** mutations cause basement membrane destabilization and tissue dysfunction in humans, nematodes, flies, and mice (Borchiellini et al., 1996; Gould et al., 2005; Gupta et al., 1997; Hudson et al., 2003; Pastor-Pareja & Xu, 2011; Pöschl et al., 2004; Rodriguez et al., 1996), whereas, Col-IV**^α345^** mutations cause a dysfunctional kidney ultrafilter in patients with Alport syndrome (Groopman et al., 2019; Hudson et al., 2003; Pokidysheva et al., 2021; Savige et al., 2023). The ubiquitous expression, evolutionary conservation and pathogenic mutations illuminate the essentiality of the Col-IV**^α121^** scaffold in animal evolution (Fidler et al., 2017, 2018; P. S. Page-McCaw et al., 2025).

In the assembly of the Col-IV**^α121^** scaffold, sulfilimine bonds crosslink Met^93^ and Hyl^211^ residues that covalently weld together the trimeric NC1-domains of adjoining triple-helical protomers, forming a globular hexamer structure at the interface (Boudko et al., 2021; R. Vanacore et al., 2009). Crosslink formation involves two halogens in their ionic forms, Cl in the initial step of hexamer assembly and Br in the final step, as a cofactor in peroxidasin-catalyzed formation of the sufilimine bond (Brown et al., 2018) (Fig, 13). The analogous mechanism is operative in Col-IV**^α345^** assembly (Pedchenko et al., 2021). Our findings in the present study reveal that bond occurrence and mechanism of formation in the Col-IV**^α121^** scaffold are conserved in a basal cnidarian (Fig. 9 and 13). Others have found that experimental reduction of bond formation in flies, nematodes, zebrafish and mice causes basement membrane dysfunction (Bhave et al., 2012; Fidler et al., 2014; Gotenstein et al., 2018; McCall et al., 2014; A. Page-McCaw & Ferrell, 2025). Collectively, these findings reveal the essentiality of the sulflimine bond in Col-IV scaffolds, enabling multicellularity, evolution and adaptation of metazoans. The essentiality posits two fundamental questions: how do sulfilimine bonds function in scaffolds and how do they enable biological function?

**Figure 13.**
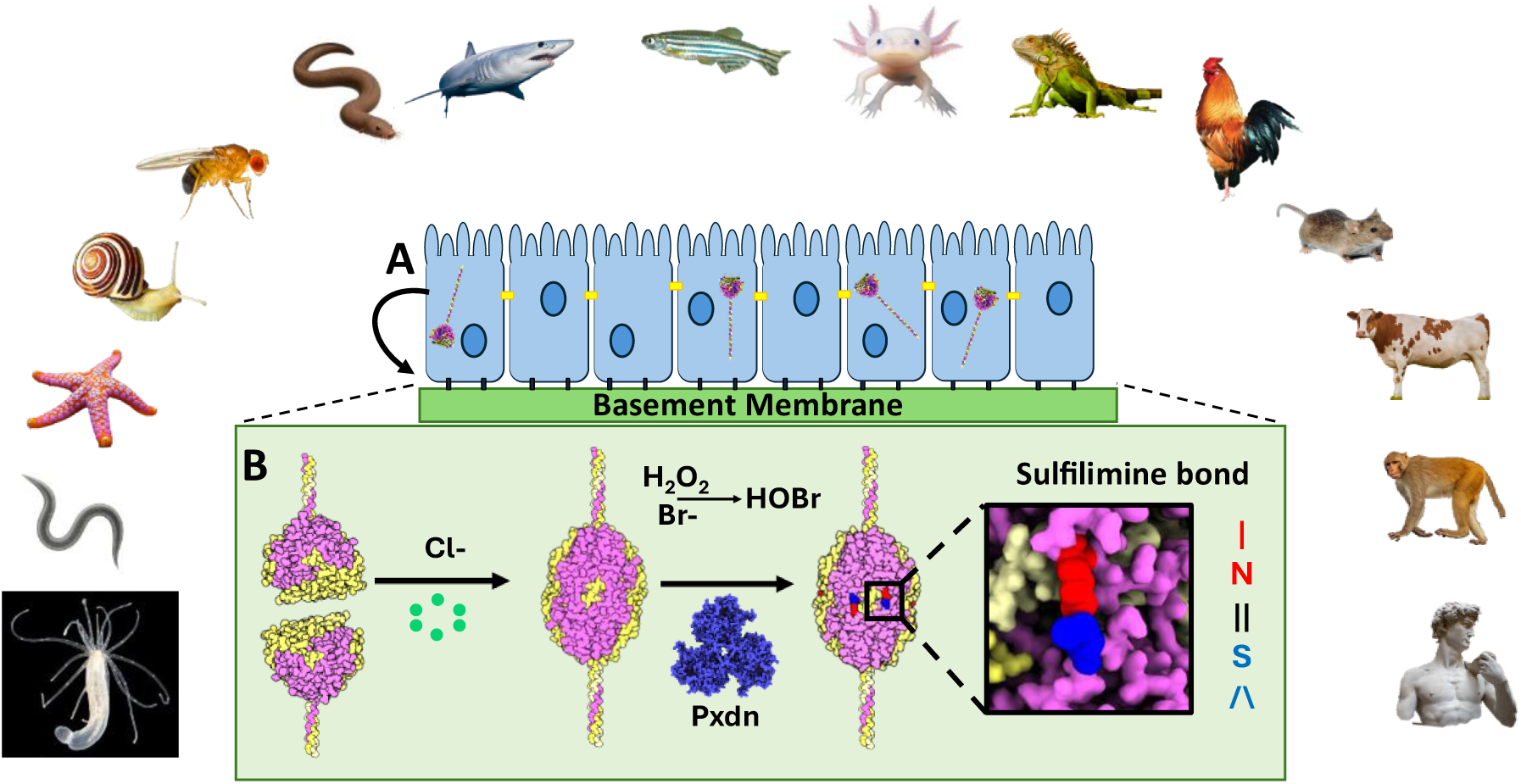
Chloride and bromine-mediated sulfilimine bonds stabilize NC1 hexamers of Col-IV^α121^ scaffold in the basement membrane across the animal kingdom. (A) Scaffold assembly begins inside the cell with the assembly of Col-IV trimeric protomers, stabilized by, among other factors, disulfide bonds. (B) Once secreted, scaffold assembly continues in the extracellular space. Here the NC1 domains of two protomers associate into a hexamer, driven by chloride pressure. Next these hexamers are covalently linked by sulfilimine bonds, driven by the enzyme peroxidasin (Pxdn) with the cofactors hydrogen peroxide and bromide that form hydrobromic acid.

In the present study of the Col-IV**^α121^** scaffold, we found that sulfilimine bonds function as a covalent reinforcement of the hexamer quaternary structure, imposed by Cl-conformational constraints (Figs. 10 and 14). Up to six sulflilimine bonds, located around the circumference and at the trimer-trimer interface, covalently interlock via a clasp-motif at the trimer-trimer interface. The interlock increases the avidity of binding interactions across all six hexamer subunits by: a) covalently linking the two trimers, b) stabilizing the β-sheet motifs in the β-pinwheel structures of trimers and stabilizing the side-by-side association of neighboring trimer subunits. Together, these features strengthen the hexamer quaternary structure, preventing trimer-trimer dissociation and side-by-side dissociation of dimer subunits (Fig. 14). Possibly these features also confer conformational constraints at the hexamer surface that are critical for interactions with binding partners (Pokidysheva et al., 2025) and for the intertwining of triple helical supra-structures within scaffolds. Moreover, hexamers are a potential structural weak point in Col-IV scaffolds, vulnerable to mechanical stress, chemical damage and proteolytic degradation (Fig. 14), whereas the triple-helical domain is highly resistant to proteases with a lifetime of millions of years (Tuinstra et al., 2025; Yang et al., 2024). In a recent study, loss of sufilimine crosslinks in mutant flies caused decreased stiffness of the Col-IV**^α121^** scaffold and tissue dysfunction (A. Page-McCaw & Ferrell, 2025). Stiffness may be a critical property associated with hexamer stability, conferred partly by sulfilimine crosslinks, that enables the tethering of laminin and other macromolecules and growth factors to scaffolds enabling cell signaling and behavior.

**Figure 14.**
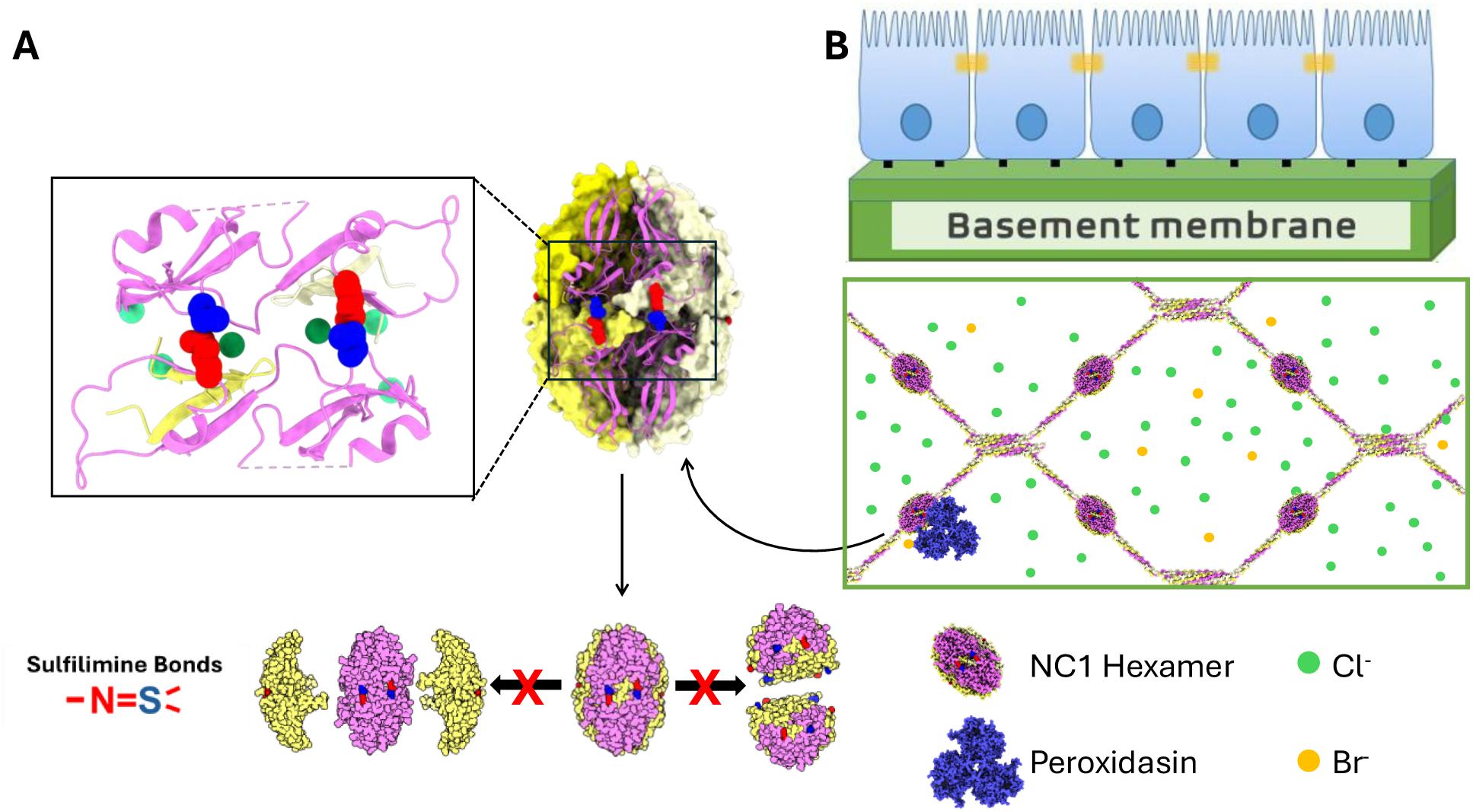
Sulfilimine bonds function as a covalent reinforcement of the hexamer quaternary structure Col-IV^α121^ scaffold of basement membranes, enabling epithelia tissue genesis, function and evolution of Animalia. (A) The NC1 hexamer is stabilized by a “clasp motif” that is governed by multiple interactions across 4 of the hexamer subunits per motif, and the sulfilimine bond covalently links two of alpha chains. The resulting hexamer is stable, and even in the absence of chloride remains in the hexametric state (B) This covalent linkage of the NC1 hexamer by the enzyme peroxidasin results in a very stable and strong Col-IV scaffold that provides a basis for formation of the basement membrane. There are also lateral interactions of collagen triple helices which are not represented in the schematic (Yurchenco & Ruben, 1987).

The essentiality of the unique sulfilimine crosslink as a reinforcement of Col-IV scaffolds posits the question of why nature utilized the sulfilimine bond rather than the common disulfide bond, both of which can be cleaved by reducing agents. A stepwise process of scaffold assembly emerged with cnidarians and conserved across the animal kingdom whereby disulfide bonds stabilize protomer assembly within the cell and sulfilimine bonds stabilize protomer oligomerization and scaffold assembly on the outside of cells. In the first step, six disulfide bonds interlock the tertiary structure of the NC1 domain of each Col-IV a chain, which in turn associate forming the trimeric NC1-domain of a triple-helical protomer. Subsequently, protomers are exported to the extracellular microenvironment of epithelial cells, where they oligomerize via their trimeric NC1-domains forming NC1-hexamers, stabilized by 36 disulfide bonds, that connect the interface of adjoining protomers (Figure 15A). Oligomerization is triggered by the extracellular Cl- gradient, which initiates and stabilizes hexamer formation and, therefore, juxtaposes Met^93^ and Hyl^211^ residues as the substrate for bond formation. Subsequently, 6 sulfilimine bonds, formed by a triad of peroxidasin, Br- and H_2_O_2_, crosslink these residues that reinforce the quaternary hexamer structure. The assembled scaffold harbors multiple binding sites that spatially and temporally organize extracellular molecules of basement membranes (Brown et al., 2017; Fidler et al., 2018; Parkin et al., 2011).

**Figure 15.**
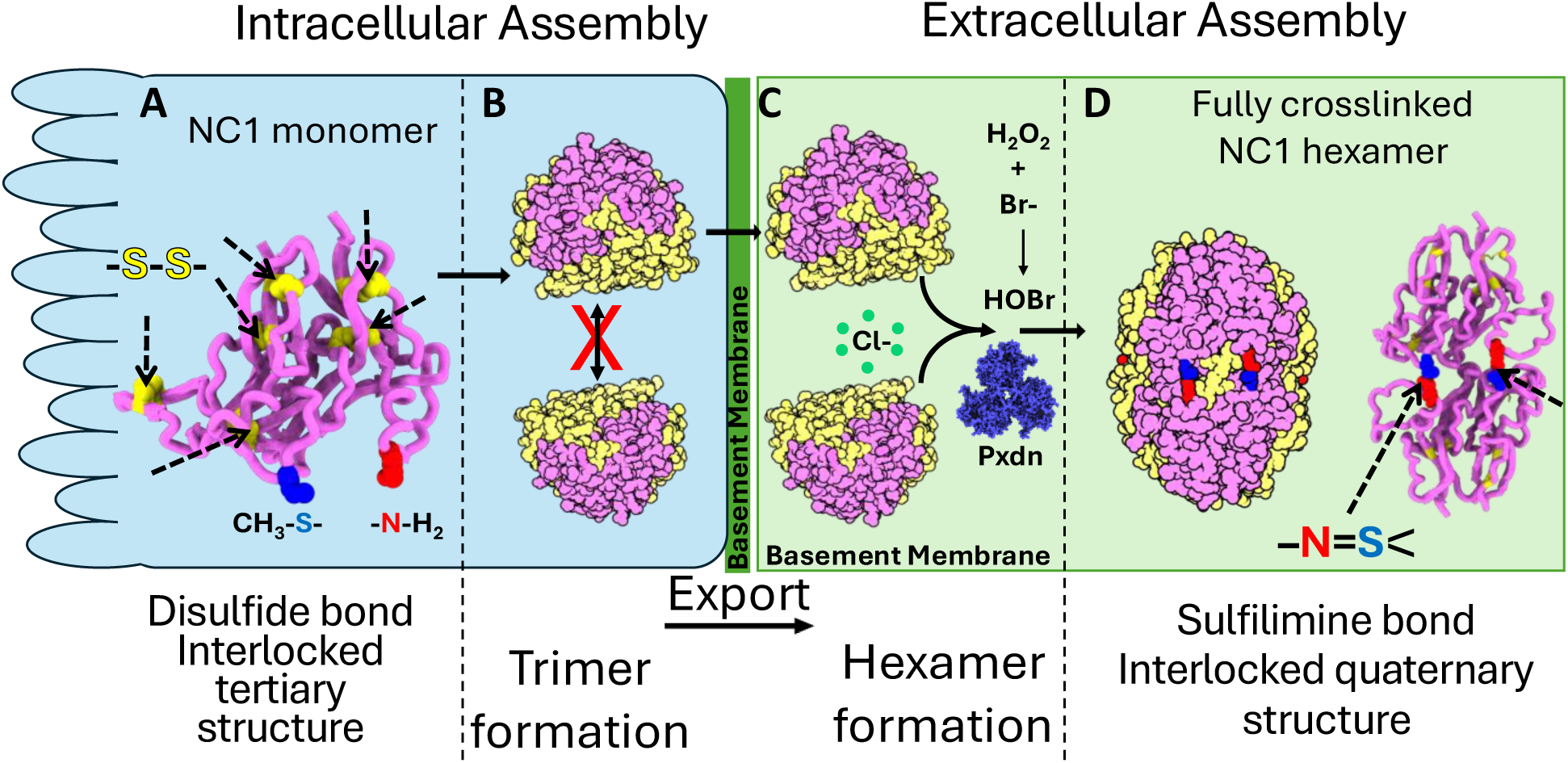
Disulfide bonds are formed within the cell, and stabilize the tertiary structure within a single NC1 monomer, whereas sulfilimine bonds are formed in the extracellular space and stabilize the quaternary structure of the NC1 hexamer. (A) Disulfide bonds are formed early in the scaffold assembly process, within the endoplasmic reticulum inside the cell, and they act to form crosslinks within a single NC1 monomer, stabilizing its tertiary structure. The assembled trimers do not form hexamers while still inside the cell, but only after they are exported to the extracellular space. (B) Hexamers are not formed while still in the secretory compartment within the cell. (C) After export, chloride pressure drives the assembly of the hexamer. Next, peroxidasin, along with HOBr forms the sulfilimine bond that crosslinks the NC1 hexamer, joining together pairs of NC1 monomers and stabilizing the quaternary structure of the hexamer (D). The spatial separation of the machinery required for formation of the sulflimine bond from the compartment where trimers are formed prevents premature assembly of the scaffold within the cell. If disulfide bonds were responsible for the stability of the quaternary structure of the hexamer, there would be no spatial separation to prevent premature assembly.

Conceivably, if a disulfide bond had been used instead of a sulfilimine bond, a seventh disulfide bond could form intracellularly within a single monomer between the residue at positions 93 and 211 (labeled as blue S and red N in Fig 15A). In this case, the NC1-monomer could assemble into an unproductive trimer and/or cause the aberrant oligomerization of protomers within the cell (Fig. 15B), hindering the assembly of a functional scaffold on the outside of cells. In contrast, the formation of sulfilimine bonds is a specific extracellular post-translational event that reinforces the assembled Col-IV scaffold. Moreover, the enzymatic mechanism, involving peroxidasin and Br- as cofactor, enables tissue specific regulation of bond formation, as exemplified by the LBM with low levels of sulfilimine bonds (Fig. 2B) as compared to the PBM which is highly crosslinked (Fig. 2E). Tissue-specific modulation may be critical to Col-IV function in various basement membranes that underlie epithelial tissues (De Gregorio et al., 2025; Sekiguchi & Yamada, 2018).

In summary, Col-IV scaffolds, a primordial and ubiquitous component of basement membranes, are reinforced during assembly by the formation of the sulfilimine bond that arose in basal cnidarians. The assembly of Col-IV scaffolds has emerged as an intricately controlled pathway requiring the use of two kinds of reducible bonds: the disulfide bond, formed intracellularly, that stabilizes tertiary structure NC1-domains of protomers; and the sulfilimine bond, formed extracellularly, that reinforces the stability of the hexamer stability of scaffolds. While the disulfide bond is common feature of protein structures, the only known occurrence of the sulfilimine bond in a biomolecule is in the NC1-hexamer of Col-IV scaffolds. Our discovery of the biological significance of the sulfiimine bond (Fidler et al., 2014; McCall et al., 2014; R. Vanacore et al., 2009) has drawn attention to sulfilimines as a class sulfur derived compounds with broad applications in organic chemistry (Arvidsson, 2025; Chen et al., 2023; Greenwood et al., 2025; Liang et al., 2023; Roy & Gauld, 2023).

## Materials and Methods

### Purification of NC1

NC1 was isolated from Bovine Lens Capsule and Placenta (from Pel-Freeze biologicals). Click or tap here to enter text.Briefly, tissues were homogenized using a Polytron in Homogenization buffer consisting of 50 mM Tris–HCl, pH 7.4, 150 mM NaCl, 20 mM EDTA and protease inhibitors. After centrifugation at 10kxg at 4°C for 30min the supernatants were discarded, and the pellets were homogenized again. This cycle was repeated for a total of 3 times. Next the pellet was homogenized in 10 mM Tris–HCl, pH 7.5, 1% deoxycholate and incubated with mixing at room temperature for 1-2hrs followed by centrifugation at 12kxg for 30min at room temperature and the supernatant was discarded. This pellet was resuspended in cold 25 mM HEPES pH 7.5, 150 mM NaCl, 25 mM 6-benzamidine–HC and centrifuged 12kxg at 4°C for 30min and the supernatant was discarded, and this was repeated for a total of 3 times. The final pellet was resuspended in the same buffer with 10 mM CaCl_2_ and 6 μg/ml collagenase (from Worthington biochemical corporation) added, and the sample was incubated with agitation at 37°C overnight. After digestion. The sample was centrifuged 12kxg at room temperature for 30min, and the supernatant containing the NC1 domain was used for analysis. NC1 was also isolated from Peroxidasin KO mice described in (Bhave et al., 2017), and purification was conducted as described previously (Boudko et al., 2018, 2023).

### Size exclusion Chromatography of NC1

Digested NC1 samples were analyzed by size exclusion chromatography using a Superdex 200 increase 10/300 GL gel filtration column (GE healthcare). Samples were injected with a 0.5ml sample loop and run at 0.5ml/minute and elution was monitored by absorbance at 280nm. Chloride containing buffer was composed of 25 mM Tris–HCl, pH 7.5, 150 mM NaCl, and Chloride free buffer was composed of 25 mM Tris-acetate, pH 7.5, with 150 mM sodium acetate.

### Cell culture and native electrophoresis

PFHR9 cells that produce a basement membrane collagen IV network with sulfilimine bonds were grown in culture as described (Bhave et al., 2012), treated with 50 µM phloroglucinol, and the NC1 fraction was collected as described above.

### N. vectensis culture

Adult *Nematostella vectensis* were cultured in the laboratory and bred using methods adapted from Darling et al. (Darling et al., 2005) and described in Tucker et al. (Tucker et al., 2011). Animals were maintained in glass dishes containing 1 μm-filtered seawater (Lake Product Company LLC, formula ASTM D 1141-52), diluted 2.5 times with distilled water to a salinity of 14%. Animals were kept at room temperature and fed *Artemia* nauplii. Water was changed twice a week before feeding. Fertilized eggs of *Nematostella vectensis* were hatched in regular media, and at the planula stage 60 animals were put in artificial media supplemented with various concentrations of bromide (normal: 220 µM, low: 0.9 µM, and bromide free) for 7 months. All animals were fed with brine shrimps washed and soaked in bromide-free medium for 1 hour to minimize food intake of bromide.

### Br-free Salt Purification

To supplement artificial culturing media with various concentrations of bromide, bromide free NaCl was purified as described previously (McCall et al., 2014). Briefly, concentrated solutions of sodium and potassium hydroxide and reagent-grade HCl were placed in a sealed chamber that prevented liquid mixing yet allowed HCl vapor to diffuse from the concentrated HCl solution to the hydroxide solutions. After 4 days, sufficient HCl vapor had diffused and neutralized the hydroxide, and the resultant chloride salt was assayed for purity via ICP-MS.

### Purification and detection of *N. vectensis* collagen IV NC1 hexamer

One hundred fifty *Nematostella* animals per treatment were collected, homogenized, and digested with bacterial collagenase as described above. Bromide free buffers were used as appropriate in all isolation and purification steps. Frozen animal pellets were sonicated in DOC Lysis Buffer (10 mM Tris pH 7.5, 1.0% deoxycholate, 0.5 mM EDTA). Insoluble material was isolated by centrifugation at 20,000×g for 20 minutes. The pellet was then washed with High Salt Buffer (1M NaCl 50 mM Tris-Cl, pH 7.5) or in Urea Buffer (2M Urea 50 mM Tris-Cl, pH 7.5) in some experiments and basement membranes were collected by centrifugation as above. Basement membrane was then washed with No Salt Buffer (10 mM Tris-Cl, pH 7.5) and digested in Collagenase Buffer (50 mM Tris-Cl pH 7.5, 5 mM CaCl_2_, 5 mM benzamidine, 25 mM 6-aminocaproic acid, 0.4 mM PMSF, 0.1 mg/mL bacterial collagenase (Worthington biochemical company)). NC1 hexamers from the supernatant of the collagenase digest were purified by gel filtration chromatography on a Superdex S200 column. Purified NC1 hexamers were then analyzed by SDS-PAGE, stained with SYPRO Ruby or Coomassie Blue. Densitometry analysis was performed (BioRad Imager).

### Western Blot detection of NC1

For western blotting detection of NC1 collagenase digested samples were resolved on SDS PAGE and transferred to nitrocellulose membranes using a transblot turbo and the standard protocol. After blocking in 5% milk TBS with 0.1%Tween, blots were developed with NC1 specific JK-2 rat monoclonal antibody (Yoshikazu Sado, Shigei Medical Research Institute), and anti-Rat HRP conjugated secondary antibody and Thermo-Scientific SuperSignal West Fempto chemiluminescent substrate.

### Mass Spectrometry

NC1 domains were resolved by SDS-PAGE, excised, and “in-gel” trypsin digested as described (R. Vanacore et al., 2009). Bands were excised and subjected to in-gel trypsin digestion. Resulting peptides analyzed by a 70 minute data dependent LC-MS/MS analysis. Briefly, peptides were autosampled onto a 200 mm by 0.1 mm (Jupiter 3 micron, 300A), self-packed analytical column coupled directly to an LTQ (ThermoFisher) using a nanoelectrospray source and resolved using an aqueous to organic gradient. A series of full scans followed by 5 data-dependent tandem mass spectra (MS/MS) was collected throughout the separation. Dynamic exclusion was enabled to minimize acquisition of redundant peptide spectra. MS/MS spectra were searched via SEQUEST against a *drosophila* or *nematostella* protein database to which the mutant caveolin sequence had been appended and that also contained reversed version for each of the entries. Identifications were filtered and collated at the protein level using Scaffold (Proteome Software) (Yates et al., 1995).

### Chloride Gels

Chloride gels were run as previously described (Boudko et al., 2023). Briefly, Tris-Glycine SDS-PAGE gels, running buffer and sample buffers were supplemented by adding 150mM NaCl. Gels were run individually at 100V for 2hrs on ice, with maximum buffer and intensive stirring to avoid overheating.

### Structure prediction and Sequence Alignment

Structures were aligned and figures were generated using ChimeraX (Pettersen et al., 2021). Sequence alignments were conducted with Clustal Omega (Madeira et al., 2024), and figures were generated with JalView (Waterhouse et al., 2009). Structure prediction was conducted with AlphaFold3 server (Abramson et al., 2024).

## Author Contributions

**Bradley P. Clarke**: Conceptualization, Methodology, Investigation, Validation, Formal analysis, Visualization, Writing draft. Writing -Reviewing and Editing; **Vadim K. Pedchenko:** Conceptualization, Methodology, Investigation, Validation, Formal analysis, Visualization; **Tetyana Pedchenko:** Investigation, Data acquisition, Writing draft, Visualization**; Monica Moran:** Investigation**; Jacob Edwards:** Investigation, Data acquisition**; Kyle Vallone:** Investigation**; Carl Darris:** Conceptualization, Investigation, Validation, Formal analysis, Visualization; **Gautam Bhave:** Conceptualization, Investigation, Validation, Formal analysis, Visualization; **Patrick P. McCaw**: Investigation, Formal analysis, Visualization, Writing draft; **Julie K. Hudson:** Conceptualization, Project administration, Funding acquisition; **Sergei P. Boudko:** Conceptualization, Formal analysis, Reviewing and Editing, Funding acquisition; **Billy G. Hudson**: Conceptualization, Resources, Writing- Reviewing and Editing; Funding acquisition.

## Funding and additional information

This work was supported by the National Institutes of Health grants R01DK018381 and R01DK131101 to B. G. H. and S. P. B. The Aspirnaut students were supported by NIH grant R25DK096999 to B. G. H.

## Acknowledgements

We would like to thank Rafi Mohammed for technical assistance and mentoring of Aspirnauts.

## References

1. Abramson, J., Adler, J., Dunger, J., Evans, R., Green, T., Pritzel, A., Ronneberger, O., Willmore, L., Ballard, A. J., Bambrick, J., Bodenstein, S. W., Evans, D. A., Hung, C. C., O’Neill, M., Reiman, D., Tunyasuvunakool, K., Wu, Z., Žemgulytė, A., Arvaniti, E., … Jumper, J. M. (2024). Accurate structure prediction of biomolecular interactions with AlphaFold 3. Nature, 630(8016), 493–500. 10.1038/s41586-024-07487-w

2. Arvidsson, P. I. (2025). Sulfilimines: An Underexplored Bioisostere for Drug Design? In Journal of Medicinal Chemistry (Vol. 68, Number 4, pp. 4056–4058). American Chemical Society. 10.1021/acs.jmedchem.5c00195

3. Bergheim, B. G., Cole, A. G., Rettel, M., Stein, F., Redl, S., Hess, M. W., Ikmi, A., & Özbek, S. (2025). Molecular dynamics of the matrisome across sea anemone life history. 10.7554/eLife.105319.1

4. Bhave, G., Colon, S., & Ferrell, N. (2017). The sulfilimine cross-link of collagen IV contributes to kidney tubular basement membrane stiffness. 10.1152/ajprenal.00096.2017.-Base

5. Bhave, G., Cummings, C. F., Vanacore, R. M., Kumagai-Cresse, C., Ero-Tolliver, I. A., Rafi, M., Kang, J. S., Pedchenko, V., Fessler, L. I., Fessler, J. H., & Hudson, B. G. (2012). Peroxidasin forms sulfilimine chemical bonds using hypohalous acids in tissue genesis. Nature Chemical Biology, 8(9), 784–790. 10.1038/nchembio.1038

6. Borchiellini, C., Coulon, J., & Le Parco, Y. (1996). The function of type IV collagen during Drosophila embryogenesis. In Arch Dev Biol (Vol. 205).

7. Boudko, S. P., Ailsworth, O., Bryant, Z. K., Cole, C., Edward, J., Edwards, D. A., Farrar, S., Gallup, J., Gallup, M., Gergis, M., Holt, A., Lach, M., Leaf, E., Mahoney, F., McFarlin, M., Moran, M., Murphy, G., Myers, C., Ni, C., … Hudson, B. G. (2023). Collagen IV of basement membranes: III. Chloride pressure is a primordial innovation that drives and maintains the assembly of scaffolds. Journal of Biological Chemistry, 299(11). 10.1016/j.jbc.2023.105318

8. Boudko, S. P., Bauer, R., Chetyrkin, S. V., Ivanov, S., Smith, J., Voziyan, P. A., & Hudson, B. G. (2021). Collagen IVα345 dysfunction in glomerular basement membrane diseases. II. Crystal structure of the α345 hexamer. Journal of Biological Chemistry, *296*. 10.1016/j.jbc.2021.100591

9. Boudko, S. P., Danylevych, N., Hudson, B. G., & Pedchenko, V. K. (2018). Basement membrane collagen IV: Isolation of functional domains. In Methods in Cell Biology (Vol. 143, pp. 171–185). Academic Press Inc. 10.1016/bs.mcb.2017.08.010

10. Brown, K. L., Cummings, C. F., Vanacore, R. M., & Hudson, B. G. (2017). Building collagen IV smart scaffolds on the outside of cells. In Protein Science (Vol. 26, Number 11, pp. 2151–2161). Blackwell Publishing Ltd. 10.1002/pro.3283

11. Brown, K. L., Hudson, B. G., & Voziyan, P. A. (2018). Halogens are key cofactors in building of collagen IV scaffolds outside the cell. In Current Opinion in Nephrology and Hypertension (Vol. 27, Number 3, pp. 171–175). Lippincott Williams and Wilkins. 10.1097/MNH.0000000000000401

12. Carvalho, J. E., Burtin, M., Detournay, O., Amiel, A. R., & Röttinger, E. (2025). Optimized husbandry and targeted gene-editing for the cnidarian Nematostella vectensis. Development (Cambridge), 152(2). 10.1242/dev.204387

13. Casino, P., Gozalbo-Rovira, R., Rodríguez-Díaz, J., Banerjee, S., Boutaud, A., Rubio, V., Hudson, B. G., Saus, J., Cervera, J., & Marina, A. (2018). Structures of collagen IV globular domains: insight into associated pathologies, folding and network assembly. IUCrJ, 5(6), 765–779. 10.1107/S2052252518012459

14. Chen, Y., Fang, D. M., Huang, H. Sen, Nie, X. K., Zhang, S. Q., Cui, X., Tang, Z., & Li, G. X. (2023). Synthesis of Sulfilimines via Selective S-C Bond Formation in Water. Organic Letters, 25(12), 2134–2138. 10.1021/acs.orglett.3c00604

15. Clay, M. R., & Sherwood, D. R. (2015). Basement Membranes in the Worm: A Dynamic Scaffolding that Instructs Cellular Behaviors and Shapes Tissues. Current Topics in Membranes, 76, 337–371. 10.1016/bs.ctm.2015.08.001

16. Colon, S., & Bhave, G. (2016). Proprotein convertase processing enhances peroxidasin activity to reinforce collagen IV. Journal of Biological Chemistry, 291(46), 24009–24016. 10.1074/jbc.M116.745935

17. Corbett, K. D., Shultzaberger, R. K., & Berger, J. M. (2004). The C-terminal domain of DNA gyrase A adopts a DNA-bending-pinwheel fold. www.pnas.orgcgidoi10.1073pnas.0401595101

18. Cummings, C. F., Pedchenko, V., Brown, K. L., Colon, S., Rafi, M., Jones-Paris, C., Pokydeshava, E., Liu, M., Pastor-Pareja, J. C., Stothers, C., Ero-Tolliver, I. A., Scott McCall, A., Vanacore, R., Bhave, G., Santoro, S., Blackwell, T. S., Zent, R., Pozzi, A., & Hudson, B. G. (2016). Extracellular chloride signals collagen IV network assembly during basement membrane formation. Journal of Cell Biology, 213(4), 479–494. 10.1083/jcb.201510065

19. Darling, J. A., Reitzel, A. R., Burton, P. M., Mazza, M. E., Ryan, J. F., Sullivan, J. C., & Finnerty, J. R. (2005). Rising starlet: The starlet sea anemone, Nematostella vectensis. In BioEssays (Vol. 27, Number 2, pp. 211–221). 10.1002/bies.20181

20. De Gregorio, V., Barua, M., & Lennon, R. (2025). Collagen formation, function and role in kidney disease. In Nature Reviews Nephrology (Vol. 21, Number 3, pp. 200–215). Nature Research. 10.1038/s41581-024-00902-5

21. Ero-Tolliver, I. A., Hudson, B. G., & Bhave, G. (2015). The ancient immunoglobulin domains of peroxidasin are required to form sulfilimine cross-links in collagen IV. Journal of Biological Chemistry, 290(35), 21741–21748. 10.1074/jbc.M115.673996

22. Fidler, A. L., Boudko, S. P., Rokas, A., & Hudson, B. G. (2018). The triple helix of collagens - An ancient protein structure that enabled animal multicellularity and tissue evolution. In Journal of Cell Science (Vol. 131, Number 7). Company of Biologists Ltd. 10.1242/jcs.203950

23. Fidler, A. L., Darris, C. E., Chetyrkin, S. V., Pedchenko, V. K., Boudko, S. P., Brown, K. L., Gray Jerome, W., Hudson, J. K., Rokas, A., & Hudson, B. G. (2017). Collagen iv and basement membrane at the evolutionary dawn of metazoan tissues. ELife, 6. 10.7554/eLife.24176.001

24. Fidler, A. L., Vanacore, R. M., Chetyrkin, S. V., Pedchenko, V. K., Bhave, G., Yin, V. P., Stothers, C. L., Rose, K. L., McDonald, W. H., Clark, T. A., Borza, D. B., Steele, R. E., Ivy, M. T., Aspirnauts, T., Hudson, J. K., & Hudson, B. G. (2014). A unique covalent bond in basement membrane is a primordial innovation for tissue evolution. Proceedings of the National Academy of Sciences of the United States of America, 111(1), 331–336. 10.1073/pnas.1318499111

25. Gotenstein, J. R., Koo, C. C., Ho, T. W., & Chisholm, A. D. (2018). Genetic suppression of basement membrane defects in caenorhabditis elegans by gain of function in extracellular matrix and cell-matrix attachment genes. Genetics, 208(4), 1499–1512. 10.1534/genetics.118.300731

26. Gould, D. B., Phalan, C. F., Breedveld, B. J., van Mil, S. E., Smith, R. S., Schimenti, J. C., Aguglia, U., van der Knapp, M. S., Heutink, P., & John, S. W. M. (2005). Mutations in Col4a1 Cause Perinatal Cerebral Hemorrhage and Porencephaly. Science, 308(5725), 1164–1167. 10.1126/science.1109418

27. Greenwood, N. S., Boyer, Z. W., Ellman, J. A., & Gnamm, C. (2025). Sulfilimines from a Medicinal Chemist’s Perspective: Physicochemical and in Vitro Parameters Relevant for Drug Discovery. Journal of Medicinal Chemistry, 68(4), 4079–4100. 10.1021/acs.jmedchem.4c02714

28. Groopman, E. E., Marasa, M., Cameron-Christie, S., Petrovski, S., Aggarwal, V. S., Milo-Rasouly, H., Li, Y., Zhang, J., Nestor, J., Krithivasan, P., Lam, W. Y., Mitrotti, A., Piva, S., Kil, B. H., Chatterjee, D., Reingold, R., Bradbury, D., DiVecchia, M., Snyder, H., … Gharavi, A. G. (2019). Diagnostic Utility of Exome Sequencing for Kidney Disease. New England Journal of Medicine, 380(2), 142–151. 10.1056/nejmoa1806891

29. Gupta, M. C., Graham, P. L., & Kramer, J. M. (1997). Characterization of 1(IV) Collagen Mutations in Caenorhabditis elegans and the Effects of 1 and 2(IV) Mutations on Type IV Collagen Distribution. In The Journal of Cell Biology (Vol. 137, Number 5).

30. Hudson, B. G., Tryggvason, K., Sundaramoorthy, M., & Neilson, E. G. (2003). Alport’s Syndrome, Goodpasture’s Syndrome, and Type IV Collagen. In n engl j med (Vol. 348). www.nejm.org

31. Khoshnoodi, J., Pedchenko, V., & Hudson, B. G. (2008). Mammalian collagen IV. In Microscopy Research and Technique (Vol. 71, Number 5, pp. 357–370). Wiley-Liss Inc. 10.1002/jemt.20564

32. Kopec, K. O., & Lupas, A. N. (2013). β-Propeller Blades as Ancestral Peptides in Protein Evolution. PLoS ONE, 8(10). 10.1371/journal.pone.0077074

33. Layden, M. J., Rentzsch, F., & Röttinger, E. (2016). The rise of the starlet sea anemone Nematostella vectensis as a model system to investigate development and regeneration. In Wiley Interdisciplinary Reviews: Developmental Biology (Vol. 5, Number 4, pp. 408–428). John Wiley and Sons Inc. 10.1002/wdev.222

34. Liang, Q., Wells, L. A., Han, K., Chen, S., Kozlowski, M. C., & Jia, T. (2023). Synthesis of Sulfilimines Enabled by Copper-Catalyzed S-Arylation of Sulfenamides. Journal of the American Chemical Society, 145(11), 6310–6318. 10.1021/jacs.2c12947

35. Madeira, F., Madhusoodanan, N., Lee, J., Eusebi, A., Niewielska, A., Tivey, A. R. N., Lopez, R., & Butcher, S. (2024). The EMBL-EBI Job Dispatcher sequence analysis tools framework in 2024. Nucleic Acids Research, 52(W1), W521–W525. 10.1093/nar/gkae241

36. McCall, A. S., Cummings, C. F., Bhave, G., Vanacore, R., Page-Mccaw, A., & Hudson, B. G. (2014). Bromine is an essential trace element for assembly of collagen IV scaffolds in tissue development and architecture. Cell, 157(6), 1380–1392. 10.1016/j.cell.2014.05.009

37. Page-McCaw, A., & Ferrell, N. (2025). Basement membrane structure and function: Relating biology to mechanics. In Matrix Biology (Vol. 141, pp. 16–31). Elsevier B.V. 10.1016/j.matbio.2025.08.004

38. Page-McCaw, P. S., Pokidysheva, E. N., Darris, C. E., Chetyrkin, S., Fidler, A. L., Gallup, J., Balser, J., Curbow, C., Harris, R., Stothers, C., Wade, K., Murawala, P., Hudson, J. K., Boudko, S. P., & Hudson, B. G. (2025). Collagen IV of basement membranes: I. Origin and diversification of COL4 genes enabling metazoan multicellularity, evolution, and adaptation. Journal of Biological Chemistry, 301(5). 10.1016/j.jbc.2025.108496

39. Paix, A., Basu, S., Steenbergen, P., Singh, R., Prevedel, R., & Ikmi, A. (2023). Endogenous tagging of multiple cellular components in the sea anemone Nematostella vectensis. Proceedings of the National Academy of Sciences of the United States of America, 120(1). 10.1073/pnas.2215958120

40. Parkin, J. Des, San Antonio, J. D., Pedchenko, V., Hudson, B., Jensen, S. T., & Savige, J. (2011). Mapping structural landmarks, ligand binding sites, and missense mutations to the collagen IV heterotrimers predicts major functional domains, novel interactions, and variation in phenotypes in inherited diseases affecting basement membranes. In Human Mutation (Vol. 32, Number 2, pp. 127–143). 10.1002/humu.21401

41. Pastor-Pareja, J. C., & Xu, T. (2011). Shaping Cells and Organs in Drosophila by Opposing Roles of Fat Body-Secreted Collagen IV and Perlecan. Developmental Cell, 21(2), 245– 256. 10.1016/j.devcel.2011.06.026

42. Pedchenko, V., Bauer, R., Pokidysheva, E. N., Al-Shaer, A., Forde, N. R., Fidler, A. L., Hudson, B. G., & Boudko, S. P. (2019). A chloride ring is an ancient evolutionary innovation mediating the assembly of the collagen IV scaffold of basement membranes. Journal of Biological Chemistry, 294(20), 7968–7981. 10.1074/jbc.RA119.007426

43. Pedchenko, V., Boudko, S. P., Barber, M., Mikhailova, T., Saus, J., Harmange, J. C., & Hudson, B. G. (2021). Collagen IVα345 dysfunction in glomerular basement membrane diseases. III. A functional framework for α345 hexamer assembly. Journal of Biological Chemistry, 296. 10.1016/j.jbc.2021.100592

44. Pettersen, E. F., Goddard, T. D., Huang, C. C., Meng, E. C., Couch, G. S., Croll, T. I., Morris, J. H., & Ferrin, T. E. (2021). UCSF ChimeraX: Structure visualization for researchers, educators, and developers. Protein Science, 30(1), 70–82. 10.1002/pro.3943

45. Pokidysheva, E. N., Redhair, N., Ailsworth, O., Page-McCaw, P., Rollins-Smith, L., Jamwal, V. S., Ohta, Y., Bächinger, H. P., Murawala, P., Flajnik, M., Fogo, A. B., Abrahamson, D., Hudson, J. K., Boudko, S. P., & Hudson, B. G. (2023). Collagen IV of basement membranes: II. Emergence of collagen IVα345 enabled the assembly of a compact GBM as an ultrafilter in mammalian kidneys. Journal of Biological Chemistry, *299*(12). 10.1016/j.jbc.2023.105459

46. Pokidysheva, E. N., Seeger, H., Pedchenko, V., Chetyrkin, S., Bergmann, C., Abrahamson, D., Cui, Z. W., Delpire, E., Fervenza, F. C., Fidler, A. L., Gaspert, A., Grohmann, M., Gross, O., Haddad, G., Harris, R. C., Kashtan, C., Fogo, A. B., Kitching, A. R., Lorenzen, J. M., … Hudson, B. G. (2021). Collagen IVα345 dysfunction in glomerular basement membrane diseases. I. Discovery of a COL4A3 variant in familial Goodpasture’s and Alport diseases. Journal of Biological Chemistry, *296*. 10.1016/j.jbc.2021.100590

47. Pokidysheva, E. N., Tufa, S. F., Keene, D. R., Hudson, B. G., & Boudko, S. P. (2025). Targeted incorporation of collagen IV to the basement membrane: A step forward for developing extracellular protein therapies. Journal of Biological Chemistry, 301(7). 10.1016/j.jbc.2025.110384

48. Pöschl, E., Schlötzer-Schrehardt, U., Brachvogel, B., Saito, K., Ninomiya, Y., & Mayer, U. (2004). Collagen IV is essential for basement membrane stability but dispensable for initiation of its assembly during early development. Development, 131(7), 1619–1628. 10.1242/dev.01037

49. Pozzi, A., Yurchenco, P. D., & Iozzo, R. V. (2017). The nature and biology of basement membranes. In Matrix Biology (Vols. 57–58, pp. 1–11). Elsevier B.V. 10.1016/j.matbio.2016.12.009

50. Rodriguez, A., Zhou, Z., My, ”, Tang, L., Meller, S., Chen, J., Bellen+, H., & Kimbrell, D. A. (1996). Identification of Immune System and Response Genes, and Novel Mutations Causing Melanotic Tumor Formation in Drosophila melanogaster.

51. Ronsein, G. E., Winterbourn, C. C., Di Mascio, P., & Kettle, A. J. (2014). Cross-linking methionine and amine residues with reactive halogen species. Free Radical Biology and Medicine, 70, 278–287. 10.1016/j.freeradbiomed.2014.01.023

52. Röttinger, E. (2021). Nematostella vectensis, an emerging model for deciphering the molecular and cellular mechanisms underlying whole-body regeneration. In Cells (Vol. 10, Number 10). MDPI. 10.3390/cells10102692

53. Roy, A., & Gauld, J. W. (2023). Sulfilimine bond formation in collagen IV. Chemical Communications, 60(6), 646–657. 10.1039/d3cc05715a

54. Savige, J., Lipska-Zietkiewicz, B. S., Watson, E., Hertz, J. M., Deltas, C., Mari, F., Hilbert, P., Plevova, P., Byers, P., Cerkauskaite, A., Gregory, M., Cerkauskiene, R., Lju-Banovic, D. G., Becherucci, F., Errichiello, C., Massella, L., Aiello, V., Lennon, R., Hopkinson, L., … Flinter, F. (2023). Correction: Guidelines for Genetic Testing and Management of Alport Syndrome (Clin J Am Soc Nephrol. 2022;17(1):143–154. doi:10.2215/CJN.04230321). In Clinical Journal of the American Society of Nephrology (Vol. 18, Number 4, p. 510). American Society of Nephrology. 10.2215/CJN.04230321

55. Sekiguchi, R., & Yamada, K. M. (2018). Basement Membranes in Development and Disease. In Current Topics in Developmental Biology (Vol. 130, pp. 143–191). Academic Press Inc. 10.1016/bs.ctdb.2018.02.005

56. Summers, J. A., Yarbrough, M., Liu, M., McDonald, W. H., Hudson, B. G., Pastor-Pareja, J. C., & Boudko, S. P. (2023). Collagen IV of basement membranes: IV. Adaptive mechanism of collagen IV scaffold assembly in Drosophila. Journal of Biological Chemistry, 299(12). 10.1016/j.jbc.2023.105394

57. Sundaramoorthy, M., Meiyappan, M., Todd, P., & Hudson, B. G. (2002). Crystal structure of NC1 domains: Structural basis for type IV collagen assembly in basement membranes. Journal of Biological Chemistry, 277(34), 31142–31153. 10.1074/jbc.M201740200

58. Than, M. E., Henrich, S., Huber, R., Ries, A., Mann, K., Kü hn, K., Timpl, R., Bourenkov, G. P., Bartunik, H. D., & Bode, W. (2002). The 1.9-Å crystal structure of the noncollagenous (NC1) domain of human placenta collagen IV shows stabilization via a novel type of covalent Met-Lys cross-link. Retrieved www.rcsb.org

59. Tucker, R. P., & Adams, J. C. (2014). Adhesion networks of cnidarians: A postgenomic view. In International Review of Cell and Molecular Biology (Vol. 308, pp. 323–377). Elsevier Inc. 10.1016/B978-0-12-800097-7.00008-7

60. Tucker, R. P., Shibata, B., & Blankenship, T. N. (2011). Ultrastructure of the mesoglea of the sea anemone Nematostella vectensis (Edwardsiidae). Invertebrate Biology, 130(1), 11–24. 10.1111/j.1744-7410.2010.00219.x

61. Tuinstra, L., Thomas, B., Robinson, S., Pawlak, K., Elezi, G., Faull, K. F., & Taylor, S. (2025). Evidence for Endogenous Collagen in Edmontosaurus Fossil Bone. Analytical Chemistry. 10.1021/acs.analchem.4c03115

62. Vanacore, R., Ham, A.-J. L., Voehler, M., Sanders, C. R., Conrads, T. P., Veenstra, T. D., Sharpless, K. B., Dawson, P. E., & Hudson, B. G. (2009). A Sulfilimine Bond Identified in Collagen IV. Science, 325(5945), 1230–1234. 10.1126/science.1176811

63. Vanacore, R. M., Shanmugasundararaj, S., Friedman, D. B., Bondar, O., Hudson, B. G., & Sundaramoorthy, M. (2004). The α1.α2 network of collagen IV reinforced stabilization of the noncollagenous domain-1 by noncovalent forces and the absence of met-lys cross-links. Journal of Biological Chemistry, 279(43), 44723–44730. 10.1074/jbc.M406344200

64. Wang, X., Harris, R. E., Bayston. Laura J., & Ashe, H. L. (2008). Type IV collagens and Dpp positive and negative regulators of signaling. Fly, 2(6), 313–315. 10.1038/nature07214

65. Waterhouse, A. M., Procter, J. B., Martin, D. M. A., Clamp, M., & Barton, G. J. (2009). Jalview Version 2-A multiple sequence alignment editor and analysis workbench. Bioinformatics, 25(9), 1189–1191. 10.1093/bioinformatics/btp033

66. Yang, J., Kojasoy, V., Porter, G. J., & Raines, R. T. (2024). Pauli Exclusion by n→π* Interactions: Implications for Paleobiology. ACS Central Science. 10.1021/acscentsci.4c00971

67. Yates, J. R., Eng, J. K., Mccormack, A. L., & Schieltz, D. (1995). Method to Correlate Tandem Mass Spectra of Modified Peptides to Amino Acid Sequences in the Protein Database. In Anal. Chem (Vol. 67). https://pubs.acs.org/sharingguidelines

68. Yurchenco, P. D., & Ruben, G. C. (1987). Basement Membrane Structure In Situ: Evidence for Lateral Associations in the Type IV Collagen Network. In The Journal of Cell Biology (Vol. 105, Number 6).

