## supplemental figures for "Collagen IV of basement membrane: V. Bromide-mediated sulfilimine bonds interlock the quaternary structure of NC1-hexamer of scaffolds enabling metazoan evolution"

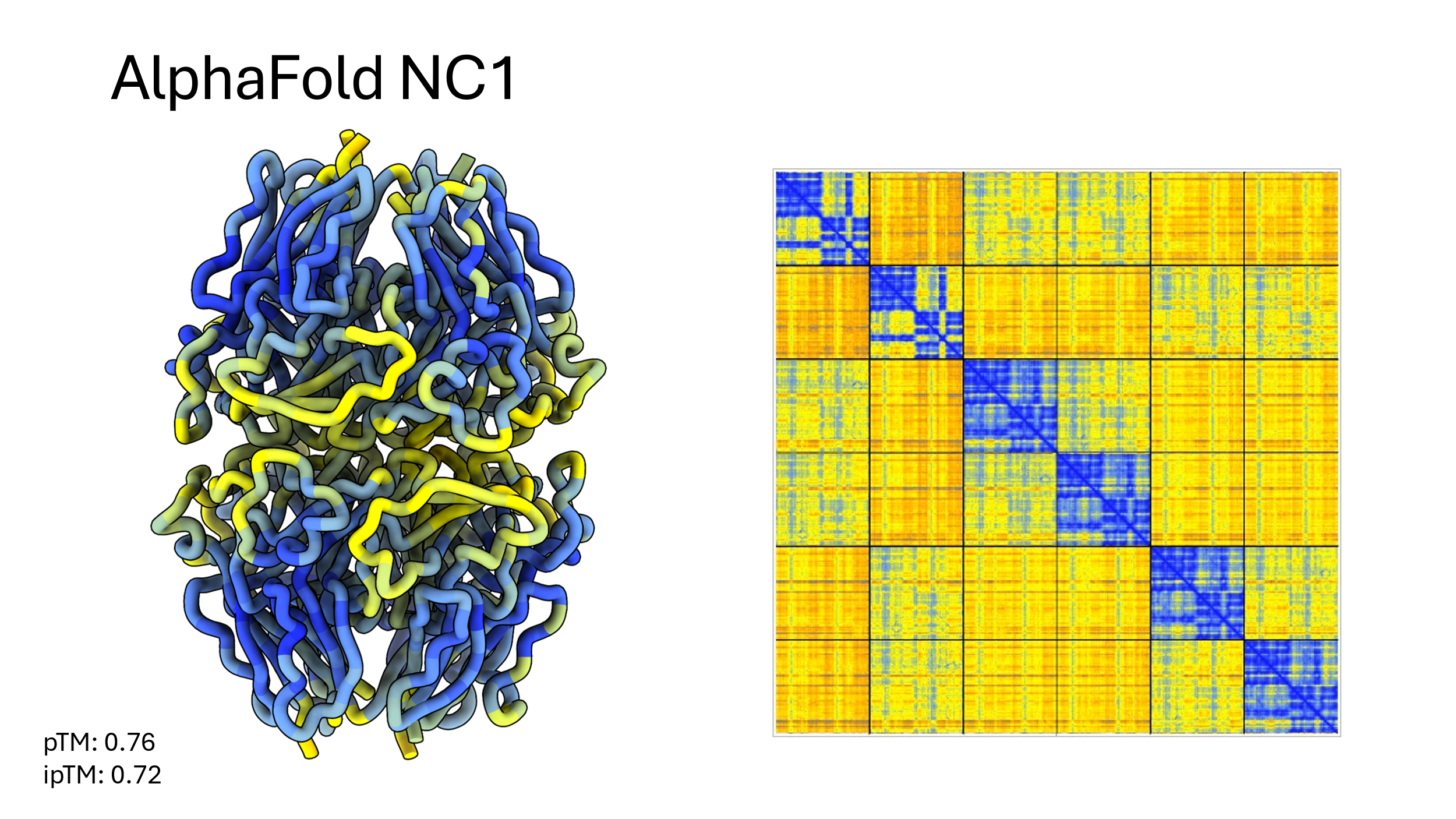

**Figure S1. AlphaFold 3 prediction of Nematostella Col-IV^α121^ NC1 hexamer**

Predicted structure of Nematostella NC1 hexamer colored by pLDDT score, and PAE plot

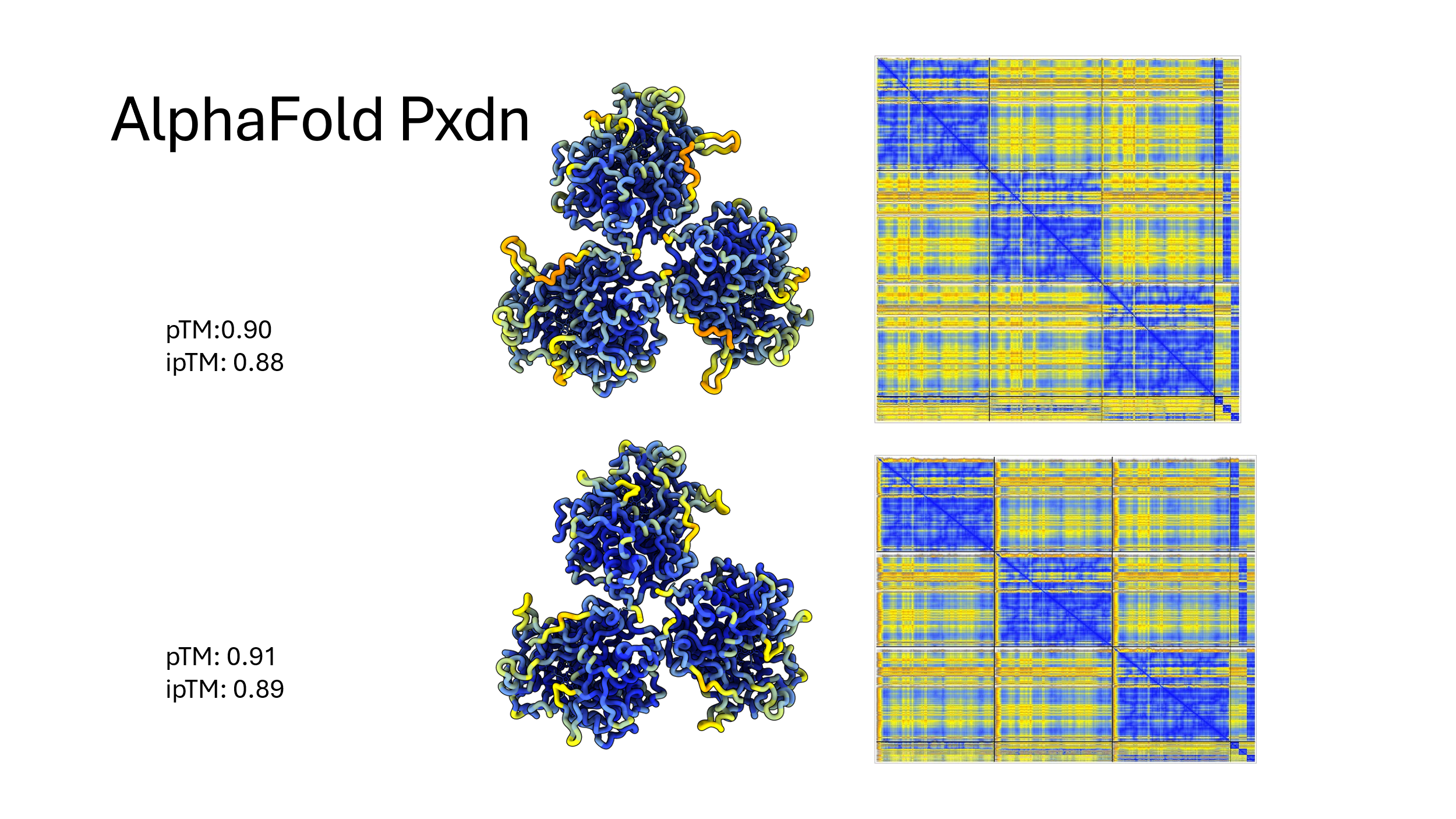

**Figure S2. AlphaFold 3 prediction of the trimeric domains of human and Nematostella peroxidasin.**

Prediction of Human (Top) and Nematostella (bottom) peroxidasin trimers colored by pLDDT score, and PAE plots

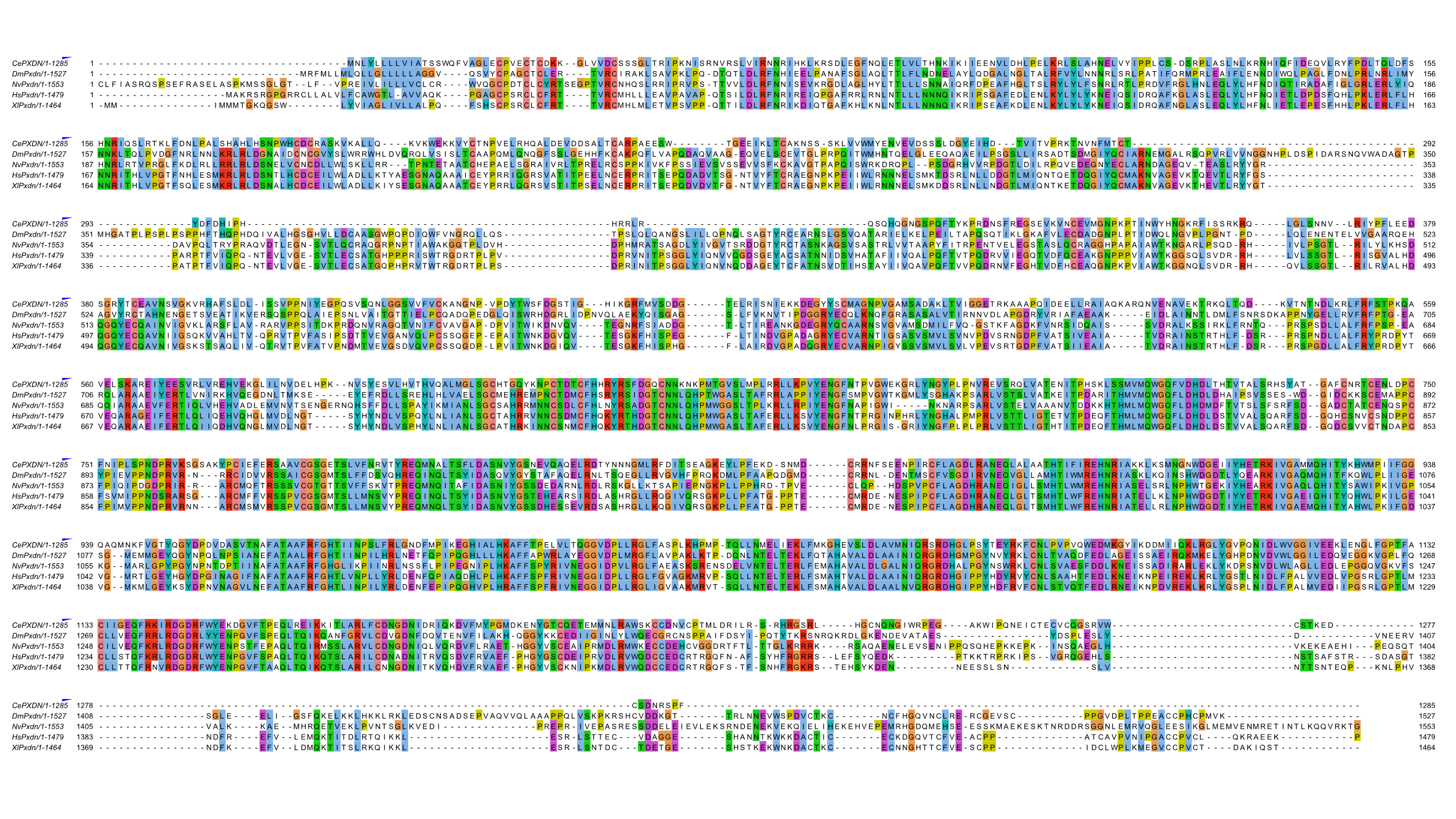

**Figure S3. Comparison of the sequence alignment of peroxidasin with other species.**

Sequence alignment of Peroxidasin from *C. elegans, D. melanogaster, N. vectensis, H. sapiens* and *X. laevis*. This alignment was used to color peroxidasin in Figure 6 B, C, E and F.

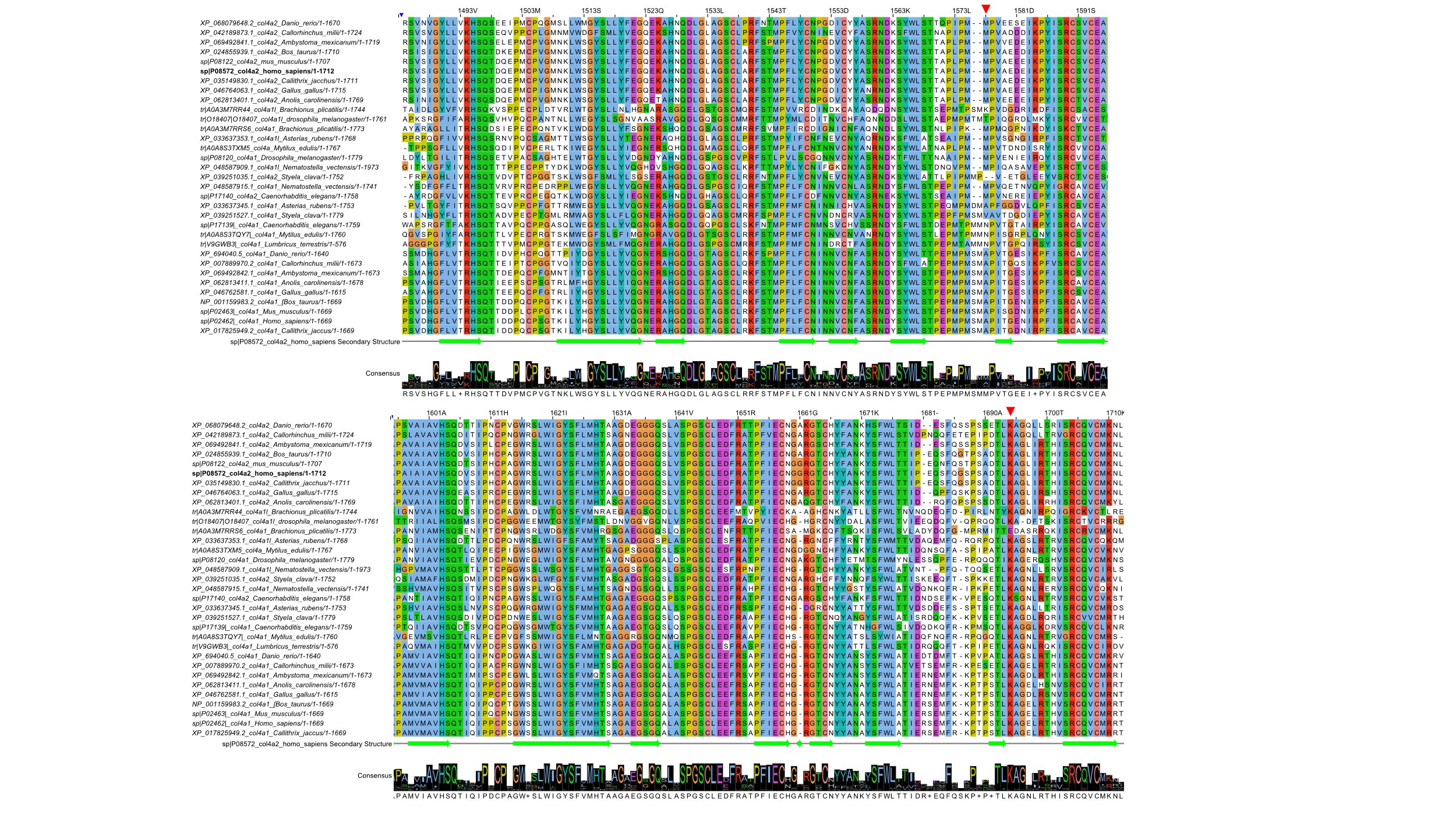

**Figure S4. Sequence alignment of Col-IV^α121^ NC1 domains.**

Sequence alignment of NC1 domains from species shown in Figure 11. Key Met and Lys residues are marked with a red arrowhead. Numbering and secondary structure (indicated below alignment) is based on *H. sapiens* Col4a2 (indicated in bold).
